# Sympathetic neuron derived NPY protects from obesity by sustaining the mural progenitors of thermogenic adipocytes

**DOI:** 10.1101/2024.05.18.594804

**Authors:** Yitao Zhu, Lu Yao, Ana Luisa Gallo-Ferraz, Bruna Bombassaro, Marcela R. Simoes, Ichitaro Abe, Jing Chen, Gitalee Sarker, Alessandro Ciccarelli, Carl Lee, Noelia Martinez-Sanchez, Michael Dustin, Kurt Anderson, Cheng Zhan, Tamas Horvath, Licio A. Velloso, Shingo Kajimura, Ana I. Domingos

**Affiliations:** Department of Physiology, Anatomy and Genetics, University of Oxford, Oxford, OX3 1PT, UK; Oxford Centre for Diabetes, Endocrinology and Metabolism Radcliffe Department of Medicine, University of Oxford, Oxford, OX3 9DU, UK; Kennedy Institute of Rheumatology, University of Oxford, Oxford, OX3 7FY, UK; Advanced Light Microscopy, The Francis Crick Institute, 1 Midland Road, London NW1 1AT, UK; Beth Israel Deaconess Medical Center, Division of Endocrinology, Diabetes & Metabolism, Harvard Medical School, Boston, MA, USA; Department of Obstetrics/Gynecology and Reproductive Sciences, Yale University School of Medicine, New Haven 06520, CT, USA; Department of Hematology, Division of Life Sciences and Medicine, University of Science and Technology of China, Hefei 230026, China; School of Sport Science, Beijing Sport University, Beijing 100084, China; Laboratory of Cell Signaling, Obesity and Comorbidities Research Center, University of Campinas, Brazil

## Abstract

Neuropeptide Y (NPY) is secreted by sympathetic nerves^1,2^, but its direct impact on thermogenic adipocytes is unknown. Here we uncover the mechanism by which peripheral NPY protects from obesity. Our imaging of cleared murine brown and white adipose tissue (BAT and WAT) established that NPY^+^ sympathetic axons are only a minority that mostly maps to the peri-vasculature; our analysis of single-cell RNA-sequencing datasets identifies mural cells as the main NPY-responsive cells in adipose tissues. We show that NPY sustains mural cells, which are known to be a source of beige cells in both BAT and WAT^3–5^ and that NPY facilitates the differentiation to thermogenic adipocytes. We found that diet-induced-obesity leads to neuropathy of NPY^+^ axons and concomitant depletion of the mural cell pool of beige fat progenitors. This defect is replicated in conditional knockout (cKO) mice with NPY specifically abrogated from sympathetic neurons. These cKO mice have whitened BAT with reduced thermogenic ability and lower energy expenditure even before the onset of obesity; they develop adult-onset obesity on a regular chow diet and are more susceptible to diet induced obesity without increasing food consumption. Our results indicate that, relative to central NPY, peripheral NPY produced by the sympathetic nerves has the opposite effect on body weight homeostasis by sustaining the proliferation of the mural cell progenitors of thermogenic adipocytes.

## Main Text

Sympathetic nerves within both brown and white adipose tissue (BAT and WAT) locally release norepinephrine (NE) to directly induce lipolysis and thermogenesis^6,7^. NPY is co-released with NE^8^, but the extent of NPY-releasing sympathetic nerves and their role in adipose tissue is not clear. Although many reports indicate that NPY in the brain stimulates appetite^9,10^, knocking out NPY has no effect on daily food consumption^11,12^, and mice without NPY receptors develop late-onset obesity despite eating less^13–15^. In humans, mutations in NPY have been linked to high body mass index (BMI) but not an unhealthy dietary pattern (https://hugeamp.org/gene.html?gene=NPY)^16^. These findings suggest that NPY from the central and sympathetic systems have contrasting impacts on maintaining body weight homeostasis. To test this hypothesis, we employed improved animal models in which NPY was selectively removed from sympathetic neurons, while leaving it intact in the brain.

## 1. In white and brown adipose tissues, NPY is exclusively sourced by one-third of TH^+^ (Tyrosine Hydroxylase-positive) sympathetic axons that preferentially innervate the vasculature

The published single-cell RNA-sequencing (scRNA-Seq) dataset of sympathetic ganglia^17^ uncovers that only two out of five clusters of sympathetic neurons express an elevated level of NPY (Supplementary Figure 1A-C). The proportion of NPY^+^ sympathetic neurons, which is about 40%, was verified by immunofluorescent co-staining of NPY and the sympathetic neuron marker tyrosine hydroxylase (TH) on superior cervical ganglia (SCG) and stellate ganglia (SG) (Supplementary Figure 1D-F). Similar TH/NPY overlap is observed on sympathetic axon bundles dissected from the inguinal adipose tissue (iWAT) (Figure 1A). We also observed that in the iWAT, NPY^+^ axons are associated with the vasculature and form plexus around the vessels (Figure 1B). To determine the distribution of TH^+^ and NPY^+^ axons in the whole adipose tissue, we performed tissue clearing of the iWAT^18^ and immunolabelled for TH, NPY and CD31, the latter of which marks the vascular endothelium. We confirmed that NPY^+^ axons are a subgroup of sympathetic axons (Figure 1C&D). By quantifying the images as in Figure 1D, we found that the double positive TH^+^NPY^+^ axons make up 32.4% of all labelled axons; 59.3% of axons are TH^+^NPY^−^ (Figure 1E-F). The images also indicate that NPY^+^ axons preferentially innervate CD31^+^ endothelium, and we confirmed that the percentage of NPY^+^ axons overlapping with CD31^+^ endothelium is higher than TH^+^ axons by quantifying images as in Figure 1D (Figure 1G). Consistent with WAT, we observed in BAT that NPY^+^ axons mainly innervate the vasculature (Figure 1H). To further determine which vessel branches are innervated by NPY^+^ axons, we quantified light-sheet images (Supplementary Figure 2A) and found that the density of NPY^+^ axons is the highest in 5^th^ order branches (Supplementary Figure 2C-D), while NPY^+^ axons do not innervate the 8^th^ order branches (Supplementary Figure 2B&D). To determine if arteries or veins are innervated by NPY^+^ axons in adipose tissue, we co-stained iWAT with NPY and EPHB4, a marker for veins^19^, or SOX17, a marker for arteries^20^. The IF images indicate that no NPY^+^ axons innervate EPHB4^+^ vessels, while they innervate SOX17^+^ vessels (Supplementary Figure2E-F). Based on these observations, we conclude that NPY^+^ sympathetic axons in adipose tissue preferentially innervate arterioles but not veins nor venules.

**Figure 1.**
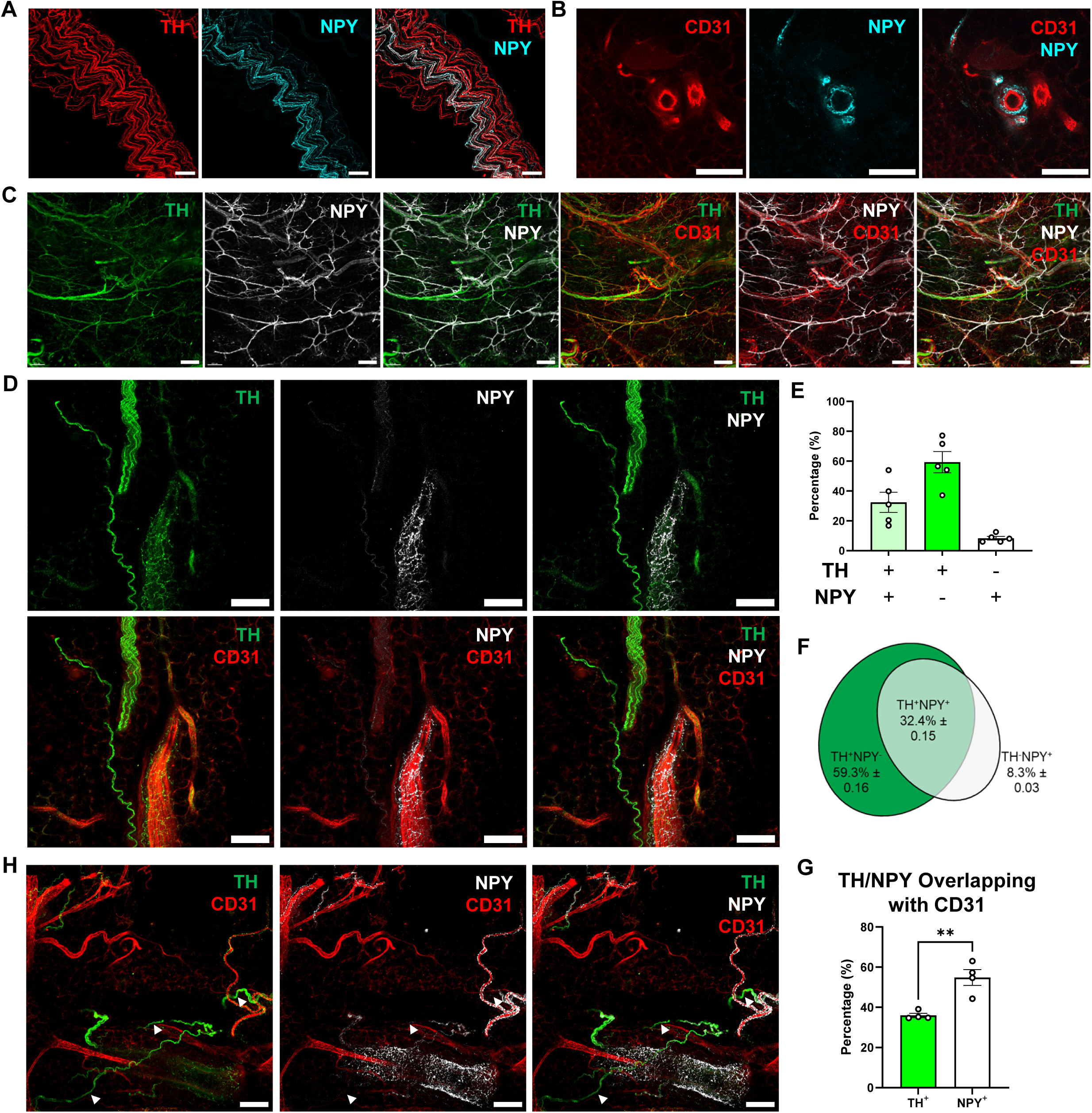
A third of TH^+^ sympathetic axons in WAT and BAT are NPY^+^, and they preferentially innervate the vasculature. (A) Confocal images of axonal bundles dissected from the iWAT of a male lean adult mouse, stained with anti-TH (red) and anti-NPY (cyan). Scale bars = 50 μm. (B) Image of a capillary within a cleared iWAT of a lean adult mouse stained with anti-CD31 (red) and anti-NPY (cyan). Scale bars=100 μm. (C) Images of cleared iWAT from lean adult male wildtype (WT) mice stained with anti-TH (green), anti-NPY (white), and anti-CD31 (red). Scale bars=50 μm. (D) Confocal images of cleared iWAT as in (C) from lean adult male mice stained with anti-TH (green), anti-NPY (white), and anti-CD31 (red). Scale bars = 100 μm. (E-F) The quantification of images as in D for TH/NPY overlap: TH^+^NPY^−^ (dark green), TH^+^NPY^+^ (light green), and TH^−^NPY^+^ (white). The percentage is shown via (E) histogram and (F) Venn diagram (n=5). (G) The quantification of images as in (D) showing the proportion of TH^+^ and NPY^+^ axons overlapping with CD31^+^ endothelial cells (n=4). (H) Confocal Images of cleared BAT from lean adult male mice stained with anti-TH (green), anti-NPY (white), and anti-CD31 (red). White arrows in the images indicate TH^+^NPY^−^ axons. Scale bars = 100 μm. All values are expressed as mean ± SEM, *p<0.05, **p<0.01, ***p<0.001, ****p<0.0001, Student T-tests.

Moreover, it is worth noting that our NPY stains are exclusively axonal. Even though earlier studies proposed that adipose-tissue macrophages (ATM) can express *Npy*^21^, this is negated by two scRNA-Seq datasets of the stromal-vascular fractions (SVFs) of murine iWAT and BAT^4,22^ (Supplementary Figure 3A-D). Consistently, by using qPCR, we do not detect expression of *Npy* on sorted ATMs from *Cx3cr1*^GFP/+^ reporter mice, despite the positivity of the *Adgre1* (F4/80) control (Supplementary Figure 3E-F).

## 2. The post-synaptic targets of NPY^+^ sympathetic neurons in WAT and BAT are NPY1R^+^ mural cells, which are progenitors of thermogenic adipocytes

We then questioned what type of perivascular cells are the postsynaptic targets of NPY^+^ axons in the adipose tissue. By reanalysing the scRNA-Seq dataset of iWAT^22^ and BAT^4^, we found *Npy1r* is highly expressed in mural cells that are marked by *Des*, *Myh11*, *Acta2* (αSMA), *Tagln*, *Cspg4* (NG2), and *Pdgfrb*, and is a bona fide marker of mural cells within WAT and BAT (Figure 2A-D; Supplementary Figure 3A-B). To validate the scRNA-Seq datasets by qPCR, we sorted mural cells (live, CD31^−^/CD45^−^/PDGFRα^−^/NG2^+^ cells) and immune cells (live, CD31^−^/CD45^+^ cells) from the SVF of iWAT and BAT (Supplementary Figure 3G) and did qPCR. Using this method, we independently confirmed that *Npy1r* and *Des* are highly expressed in mural cells but not in immune cells (Figure 2E). To confirm the existence of NPY1R in mural cells at the protein level, we dissected capillaries from adipose tissue and stained them with anti-CD31 and anti-NPY1R. We detected an NPY1R^+^ cell wrapping the capillary (Figure 2F), whose morphology matches that of mural cells of fine capillaries, which are conventionally named as pericytes^23^. To confirm the colocalization of NPY1R with multiple mural cell marker observed in scRNA-Seq datasets (Supplementary Figure 4A), we immunolabelled vessel-containing samples dissected from adipose tissues of Npy1r^Cre^; Rosa26^tdTomato^ mice for multiple mural cell markers including DES, αSMA, and NG2, and the images indicate that DES^+^, αSMA^+^, or NG2^+^ mural cells are all labelled with NPY1R-tdTomato (Figure 2G, Supplementary Figure 4B-E). Based on the observations above, we demonstrate that mural cells are the target of NPY^+^ axons and that they are NPY1R ^+^.

**Figure 2.**
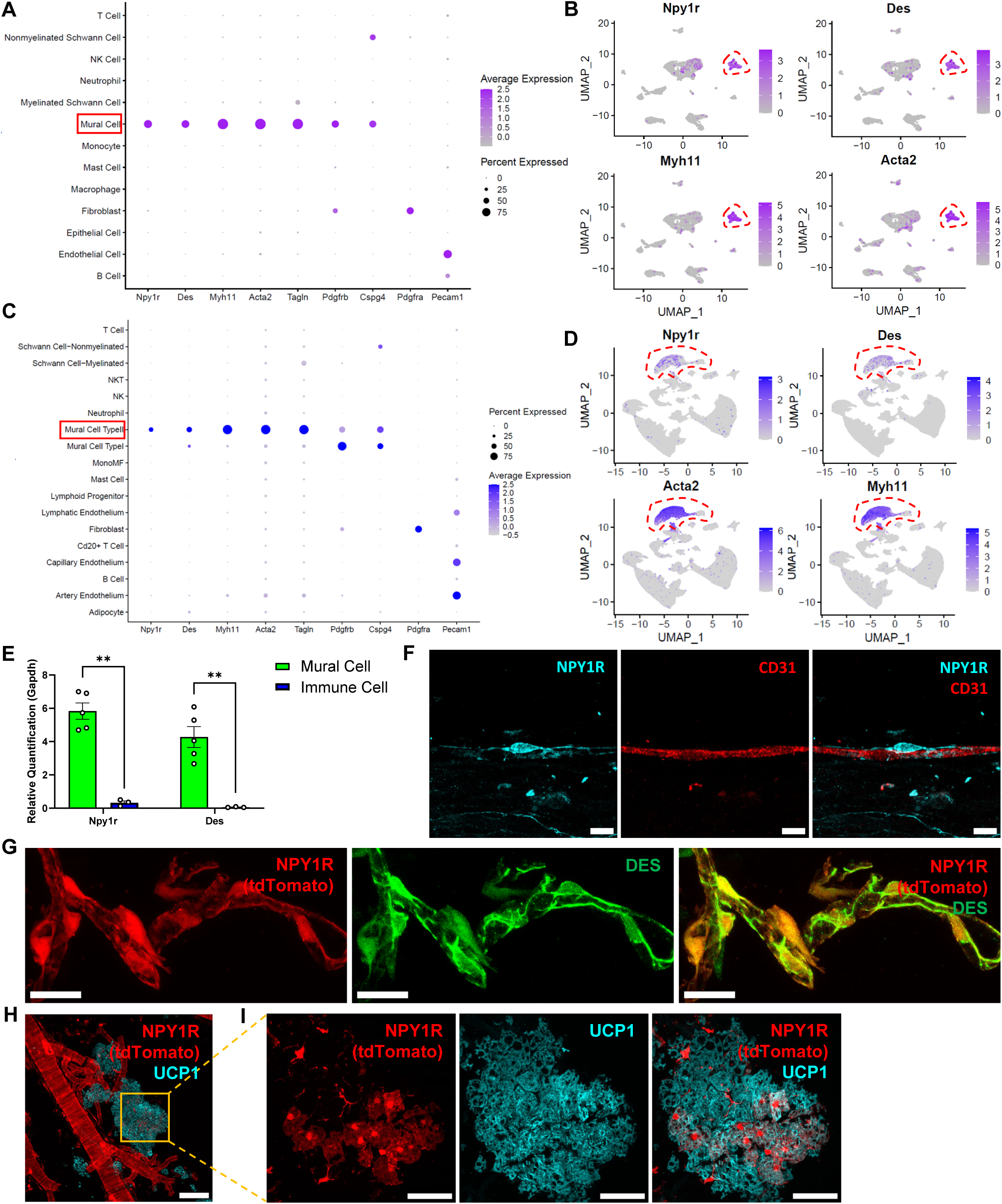
*Npy1r* is mainly expressed by mural cells, and *Npy1r*^+^ mural cells are a source of thermogenic adipocytes^3–5^. (A&C) Dot plot showing the expression of *Npy1r, Des, Myh11, Acta2 (aSMA), Tagln, Cspg4* (NG2), *Pdgfrb, Pdgfra, and Pecam1* (CD31) in the stromal vascular fraction (SVF) of (A) iWAT^22^ or (C) BAT^4^. (B&D) Embedding plots showing the expression of *Npy1r, Des, Myh11,* and *Acta2 (aSMA)* in the SVF of (B) iWAT^22^ and (D) BAT^4^. Mural cell clusters are highlighted with red rectangles or red dashed lines. (E) The expression of *Npy1r* and *Des* in sorted CD31^−^/CD45^−^/PDGFRα^−^/NG2^+^ mural cells and CD45^+^ immune cells (n=5 & 3). (F) Confocal image of the capillary dissected from the iWAT of WT mice stained with anti-NPY1R (cyan) and anti-CD31 (red). Scale bar =10 μm. (G) Confocal images of vessels dissected from Npy1r^Cre^; Rosa26^tdTomato^ mice stained with anti-DES (green), Scale bar=20 μm. (H-I) Confocal images of vessels dissected from the BAT of ND-treated, RT-housed 12-week-old Npy1r^Cre^; Rosa26^tdTomato^ mice stained with anti-UCP1 (cyan), Scale bar = 100 μm. (I) Zoomed-in images of H, scale bar = 50 μm. All values are expressed as mean ± SEM, *p<0.05, **p<0.01, ***p<0.001, ****p<0.0001, Student T-tests.

It is worth noticing that earlier reports of *Npy2r* and *Npy5r* expression by immune cells, preadipocytes and adipocytes^24–27^ are negated by our analysis of the two scRNA-Seq datasets of both BAT and iWAT, where *Npy5r* is undetectable, and *Npy2r* expression is only detectable in iWAT but negligible (Supplementary Figure 5A-B). Also, some previous reports of *Npy1r* expression^28^ by adipocytes or macrophages are negated by a single-nucleus atlas of mouse white adipose tissue and a scRNA-Seq dataset of BAT^4,29^ as well as our qPCR result of sorted ATMs (Supplementary Figure 3F). Therefore, other than mural cells, no immune cells or adipocytes are likely to be the postsynaptic targets of NPY^+^ axons in adipose tissue.

Mural cells can give rise to thermogenic adipocytes^3–5^, and previously identified progenitors of beige cells express *Npy1r* and mural cell markers^5^ (Figure 3C-E). We confirmed that mural cells are progenitors of thermogenic adipocytes by isolating mural cells from adipose tissue and differentiating them *in vitro*^30^ (Supplementary Figure 6A-B); we observed that they are more potent in differentiating into thermogenic adipocytes than stromal vascular fraction (SVF) (Supplementary Figure 6C-D). To demonstrate that NPY1R^+^ mural cells are progenitors of thermogenic adipocytes under physiological conditions, we immunolabelled BAT of Npy1r^Cre^; Rosa26^tdTomato^ room-temperature (RT) –housed mice for UCP1; using this lineage-tracing approach we observe that both mural cells and a subgroup of UCP1^+^ adipocytes are co-labelled with NPY1R-tdTomato (Figure 2H-I). Since adipocytes do not express Npy1r (Figure 2C), we reason that these NPY1R-tdTomato^+^UCP1^+^ thermogenic adipocytes are lineage-traced to NPY1R^+^ mural cells. Based on the observations above and published literature^3–5^, we conclude that NPY1R^+^ mural cells are progenitors of thermogenic adipocytes.

**Figure 3.**
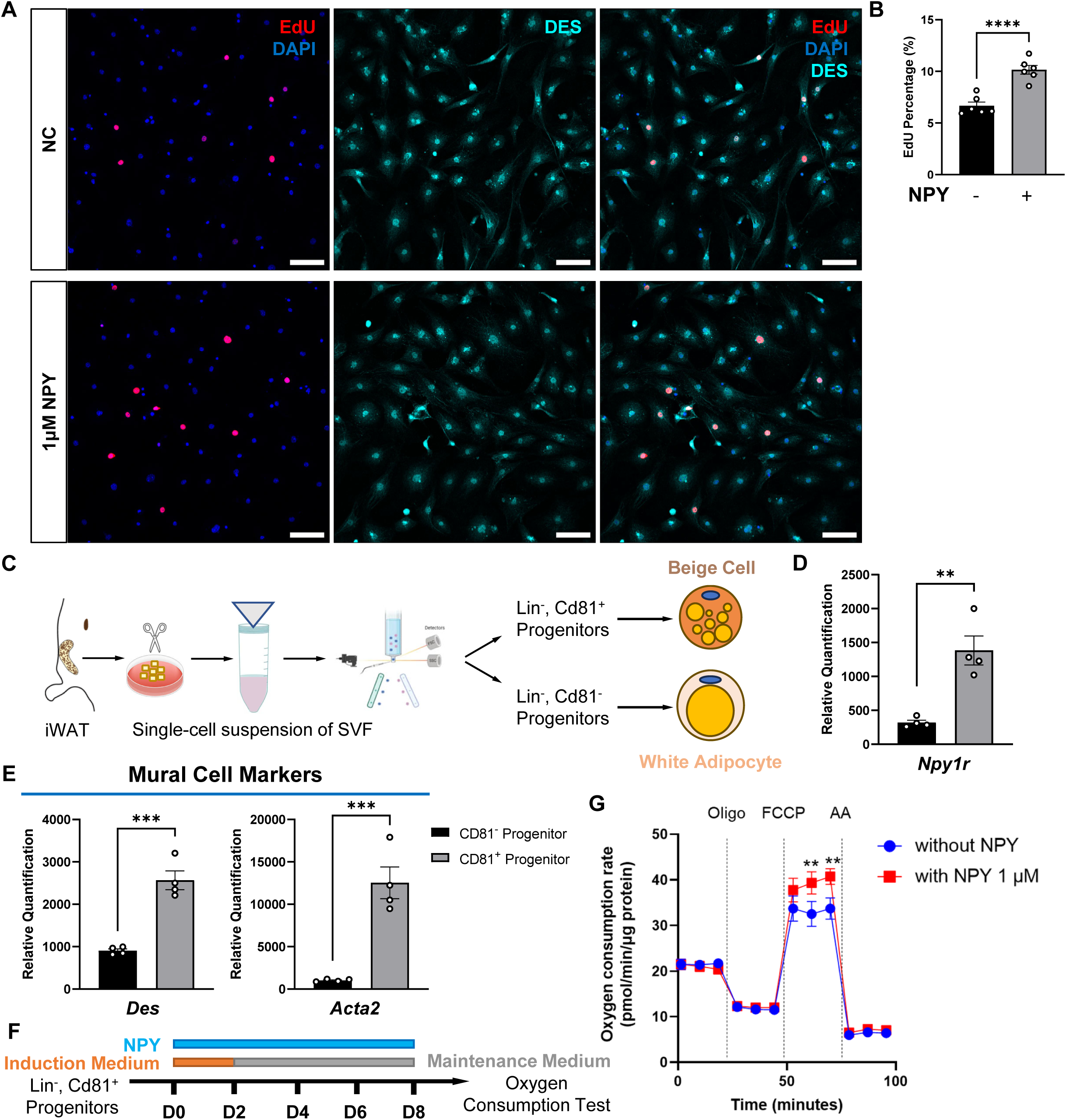
NPY promotes the proliferation of mural cells and facilitates beiging. (A) Confocal images of isolated mural cells treated with or without NPY stained with anti-DES (cyan) and DAPI (blue). EdU (red) was used to indicated proliferation. Scale bar=80 μm. (B) The percentage of EdU^+^ cells quantified based on images as in A (n=6). (C) Schematic of isolating the progenitors of thermogenic adipocytes in published paper^5^. (D-E) The expression level of (D) *Npy1r*, and (E) mural cell markers *Des* and *Acta2* in CD81^−^ progenitors of WAT and CD81^+^ progenitors of beige cells. (F) Schematic of differentiating isolated Lin^−^, CD81+ progenitors. (G) Oxygen consumption in adipocytes differentiated from CD81^+^ progenitors with or without NPY (n=5). All values are expressed as mean ± SEM, *p<0.05, **p<0.01, ***p<0.001, ****p<0.0001, Student T-tests.

## 3. NPY promotes mural cell proliferation and sustains their beiging ability

NPY1R is a kind of G protein-coupled receptor, and it can activate MAPK pathways after binding to NPY^31,32^. Since MAPK pathways a canonical regulator of cell proliferation^33^, we then questioned if NPY can promote the proliferation of mural cells. By isolating mural cells and adding NPY, we observed that NPY promotes the proliferation of DES^+^ mural cells revealed by EdU labelling (Figure 3A-B). To determine whether the effect of NPY on proliferation can affect beiging, we differentiated the progenitors of beige cells with NPY and found that the maximal respiratory capacity is increased (Figure 3F-G). Using SVF, we also demonstrate that NPY can upregulate beiging and thermogenesis genes (Supplementary Figure 6E-G). Since NPY does not affect general adipogenesis (Supplementary Figure 6H, Supplementary Figure 7B-C), as demonstrated *in vitro* using SVF and NPY1R^+^ 3T3-L1 cells (Supplementary Figure 7A), the upregulation of beiging markers is due to NPY’s effect of increasing the proportion of mural cells in the SVF, which are progenitors of thermogenic adipocytes.

Previous reports have demonstrated that mural cells can be regulated by many factors, one of which is PDGF-B^34^. On one hand, it is a factor required for the recruitment of mural cells; on the other hand, elevated levels of PDGF-B can decrease mural cells both *in vitro* and *in* vivo by upregulating fibroblast markers and promoting the proliferation of fibroblasts^35,36^. We then questioned whether NPY can sustain mural cells when being challenged with PDGF-B. This is physiologically relevant since diet-induced-obesity (DIO) increases PDGF-B in adipose tissue^36^. By treating isolated mural cells with PDGF-BB, we found that PDGF-BB indeed decreased the proportion of mural cells and increased the number of spindle-shaped PDGFRα^+^ fibroblasts, an effect that is counteracted by NPY (Supplementary Figure 8A-B). Furthermore, using qPCR, we demonstrate that NPY can restore the expression of *Npy1r* and other mural cell markers including *Pdgfrb*, *Rgs5*, and *Angpt1* that are downregulated by PDGF-BB (Supplementary Figure 8C-E). Since mural cells have been identified as the progenitor of thermogenic adipocytes, we then examined whether NPY can preserve their beiging ability against PDGF-BB. We observe that PDGF-BB-treatment downregulates thermogenesis genes and beiging genes, while NPY can restore their expression (Supplementary Figure 8F-G). Based on the observations above, we demonstrate *in vitro* that NPY can sustain mural cells as the progenitors of thermogenic adipocytes

## 4. High-fat diet-induced obesity depletes NPY+ innervation and mural cells in adipose tissue, increasing vascular leakiness

To test whether NPY^+^ sympathetic axons are also affected by the obesity-induced sympathetic neuropathy in the adipose tissue^37^, we immunolabelled the cleared iWAT of normal diet (ND)– and high-fat diet (HFD)-treated mice for NPY and CD31. Our light-sheet images show that the density of NPY^+^ axons is significantly decreased in the iWAT of HFD-treated mice compared with ND-treated lean mice and, as a result, the percentage of vessels innervated by NPY^+^ axons is significantly decreased as well (Supplementary Figure 9A, B). Consistently, NPY concentration also diminished in the iWAT of HFD-induced obese mice and leptin-deficient obese mice (Supplementary Figure 9C). We can ascertain that this is not due to any changes in *Npy* expression in the ganglionic neuronal soma (Supplementary Figure 10A-B) or systemic NPY level (Supplementary Figure 9D). Altogether, these results show that obesity-induced sympathetic neuropathy reduces the NPY^+^ innervation in the adipose tissue, and this is sufficient to decrease the local concentration of NPY in the iWAT. We then asked if mural cells, as postsynaptic targets of NPY^+^ innervation in iWAT, were also affected by obesity. By immunolabelling cleared iWATs for DES and CD31, we discovered that there were fewer mural cells covering the vasculature of the iWAT of DIO mice compared with age-matched lean mice (Supplementary Figure 9E-F). Since mural cells are needed for vascular integrity^38^, we also observed increased vascular leakiness in the iWAT of DIO mice revealed by PV-1 staining^39^ (Supplementary Figure 9G-H). To investigate if this phenomenon is related to NPY concentration, we immunolabelled sympathetic ganglia for DES to ascertain if the mural cell coverage changes when the NPY level is not affected. We indeed observed that there is no significant difference in the mural cell coverage of ganglionic vasculature between DIO mice and lean mice (Supplementary Figure 10B-E), which confirms that mural cell coverage positively correlates with the NPY level.

## 5. Loss of function of NPY from sympathetic neurons depletes mural cells and increased vascular leakiness in adipose tissue

To test the hypothesis that NPY+ innervation is required for sustaining mural cells *in vivo*, we generated a mouse model in which NPY was abrogated from sympathetic neurons: Th^Cre^; Npy^flox/flox^ mice. We first confirmed that in Th^Cre^; Npy^flox/flox^ mice, NPY is ablated from sympathetic ganglia (Supplementary Figure 11A-B) and sympathetic axons innervating iWAT and BAT (Supplementary Figure 11C&E), while general TH^+^ sympathetic innervation is not affected (Supplementary Figure 11D&F). Using this mouse model, we demonstrate that ablating NPY downregulates mural cell markers in the BAT of ND-treated mice (Figure 4A) and depletes mural cells in the BAT and iWAT of both ND-treated and HFD-treated mice (Figure 4C-D&F, Supplementary Figure 12A-B). Since mural cells are required for vascular integrity^38^, we observed upregulated pro-inflammatory genes in BAT (Figure 4B), increased vascular leakiness in iWAT (Figure 4C), and increased immune infiltration in BAT and iWAT of ND-treated Th^Cre^; Npy^flox/flox^ mice (Figure 4C&E). These observations confirmed that NPY is required for sustaining mural cells in adipose tissue and preventing immune infiltration.

**Figure 4.**
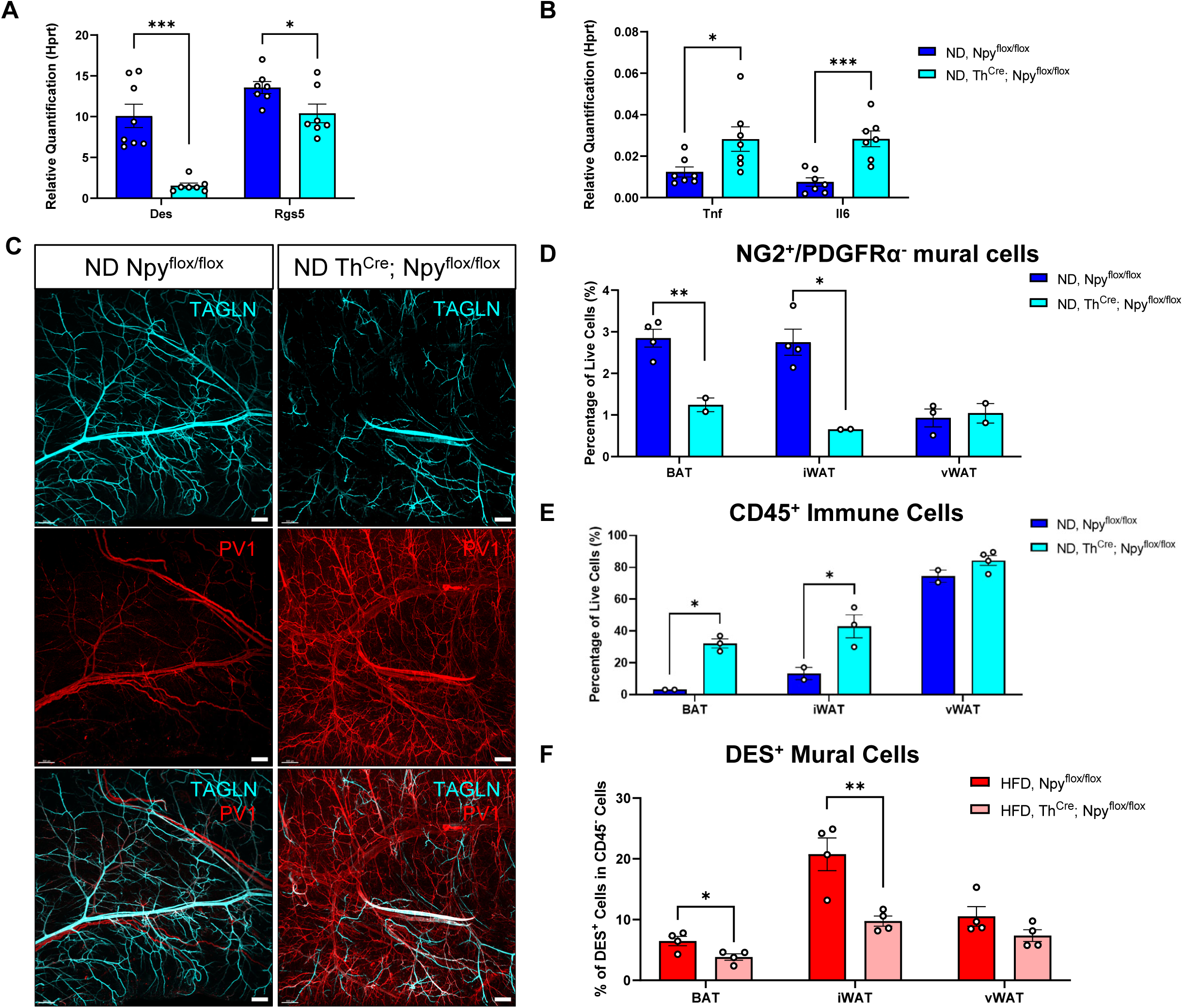
Loss of function of NPY from sympathetic neurons depletes mural cells and increases vascular leakiness. (A-B) The expression levels of (A) mural cell markers *Des* (n=8)*, Rgs5* (n=7), and (B) pro-inflammatory genes *Tnf* (n=7), and *Il6* (n=7) in the BATs of ND-treated 12-week-old male Th^Cre^; Npy^flox/flox^ mice and WT mice. (C) Light-sheet image of cleared iWAT of 30-week-old, ND-treated mice stained with anti-PV1 (red) and anti-TAGLN (cyan). Scale bar=500 μm. (D-E) The percentage of (D) PDGFRα^−^ /NG2^+^ mural cells (n=4&2) and (E) CD45^+^ immune cells (n=2&3) in BATs, iWATs, and vWATs of 30-week-old, ND-treated mice measured by flow cytometry. (F) The percentage of DES^+^ mural cells (n=4) in BATs, iWATs, and vWATs of 17-week-old, HFD-treated mice measured by flow cytometry. All values are expressed as mean ± SEM, *p<0.05, **p<0.01, ***p<0.001, ****p<0.0001, Student T-tests.

## 6. Loss of function of NPY from sympathetic neurons whitens BAT before the onset of obesity, reduces beiging, energy expenditure, and thermogenesis, and render mice more susceptible to diet-induced obesity (DIO) without increasing food intake

To test the hypothesis that NPY locally sourced from sympathetic axons in adipose tissues preserves the mural cell pool and is required for energy homeostasis, we characterised the metabolism of Th^Cre^; Npy^flox/flox^ mice. We discovered that ND-treated lean Th^Cre^; Npy^flox/flox^ mice have whitened and bigger BAT (Figure 5A&H), and their BATs are dysfunctional in thermogenesis as shown by lower levels of beiging gene *Pparg* and thermogenesis genes *Prdm16*, *Cidea*, and *Pgc1a* (Figure 5B-C). Consistent with this, Th^Cre^; Npy^flox/flox^ mice have lower daily energy expenditure (EE) (Figure 5D, Supplementary Figure 13A) and higher respiratory exchange (RER) (Supplementary Figure 13D-E) – the latter is indicative of lower usage of fat as a metabolic substrate. By food-restricting mice, we observed that Th^Cre^; Npy^flox/flox^ mice have lower BAT temperature, and they lose less weight during a 14-hour fast, which is indicative of energy conservation (Figure 5E-F). It is unlikely these alterations are related to muscular activity because Th^Cre^; Npy^flox/flox^ mice have the same lean mass and locomotor activity as WT mice (Supplementary Figure 13B-C). As a result of deficient thermogenic ability and reduced EE, ND-treated Th^Cre^; Npy^flox/flox^ mice develop adult-onset obesity at the age of 30 weeks, with heavier adipose deposit compared with WT mice (Figure 5G-J). When on an HFD, Th^Cre^; Npy^flox/flox^ mice get obese earlier and faster than WT mice, without increasing food intake (Figure 5K-L). Also, HFD-treated Th^Cre^; Npy^flox/flox^ mice have significantly heavier adipose deposit (Figure 5M-N), and their BATs contain larger lipid droplets in average (Figure 5O, Supplementary Figure 13F). Lastly, the metabolic phenotype is invariable to gender as HFD-treated female Th^Cre^; Npy^flox/flox^ mice also have heavier body weights, WATs, and BATs compared with Npy^flox/flox^ control mice (Supplementary Figure 16B-C).

**Figure 5.**
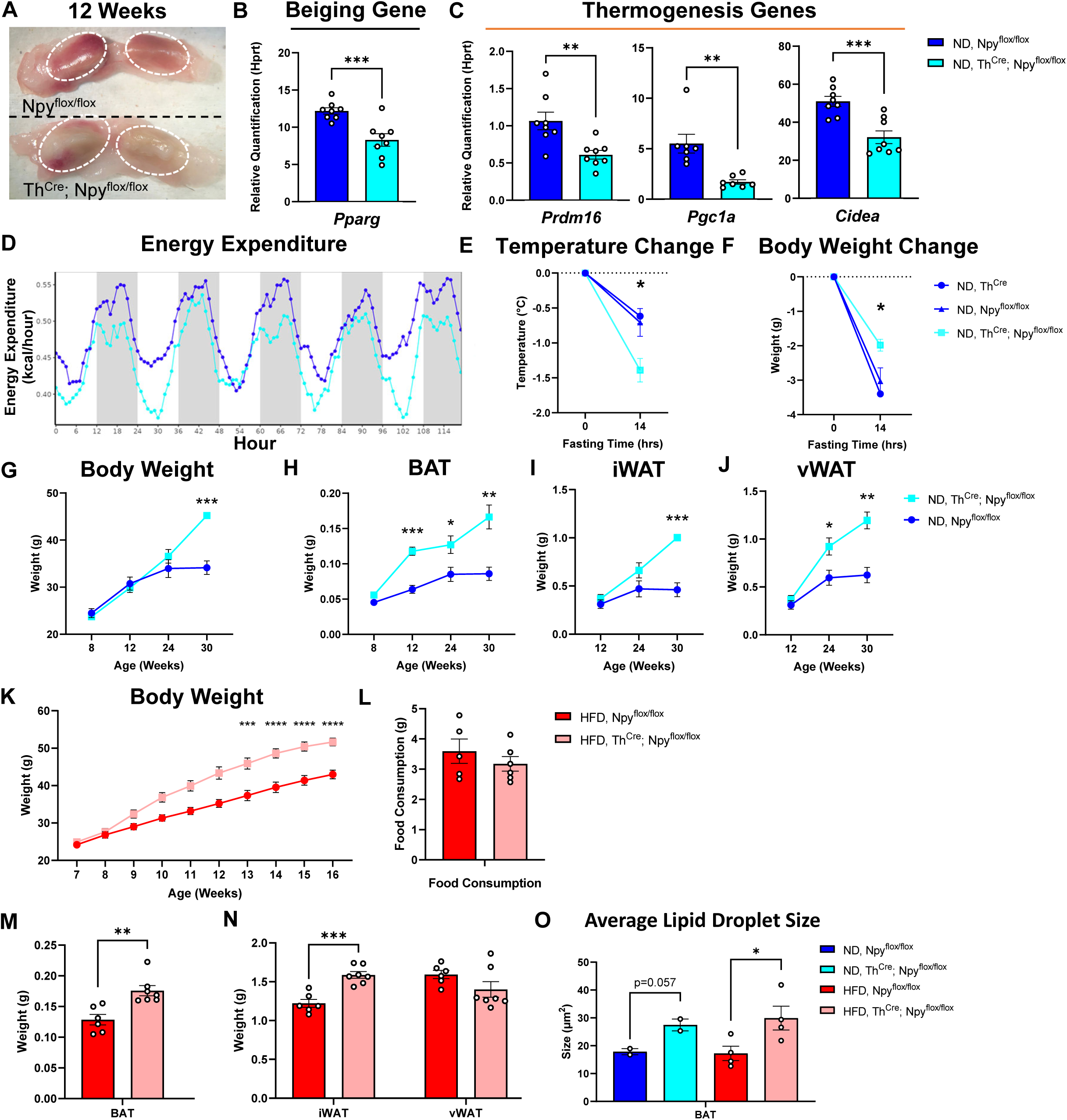
Loss of function of NPY from sympathetic neurons whitens BAT before the onset of obesity, decreases energy expenditure and thermogenesis leading to adult-onset obesity on a normal diet, and renders mice more susceptible to diet-induced obesity (DIO) without increasing food intake. (A) The picture of BATs dissected from ND-treated 12-week-old male WT (top) and Th^Cre^; Npy^flox/flox^ (bottom) mice. The dashed lines encircle the lobes of BATs. (B-C) The expression levels of (B) beiging genes *Pparg* (n=7 & 8), and (C) thermogenesis genes *Prdm16* (n=8), *Pgc1a* (n=7) and *Cidea* (n=8) in the BATs of ND-treated 12-week-old male Th^Cre^; Npy^flox/flox^ mice and WT mice (n=7). (D) Daily energy expenditure of 7-week-old male Th^Cre^; Npy^flox/flox^ mice and Npy^flox/flox^ WT mice measured using an indirect calorimetry system (n=7). (E-F) The change in (E) BAT temperature and (F) body weights of 12-week-old mice after a 14-hour fast (n=6 for Th^Cre^; Npy^flox/flox^ mice and n=3 for both Th^Cre^ and Npy^flox/flox^ WT mice). (G-J) The weights of (G) whole body, (H) BATs, (I) iWATs, and (J) vWATs of ND-treated male Th^Cre^; Npy^flox/flox^ and WT mice (Npy^flox/flox^) at different ages indicated in the plot (n=4). (K-L) (K) weekly body weights (n=8) and (L) averaged daily food consumption (n=4&6 cages) of male Th^Cre^; Npy^flox/flox^ mice and WT mice treated with HFD. (M-N) The weights of (M) BATs (n=5&6), (N) iWATs (n=6&7), and vWATs (n=6&7) of HFD-treated 17-week-old male Th^Cre^; Npy^flox/flox^ mice and WT mice. (O) The average size of lipid droplets in the BATs of ND-treated and HFD-treated male Th^Cre^; Npy^flox/flox^ mice and WT mice (n=2 for ND-treated mice, and n=4 for HFD-treated mice). All values are expressed as mean ± SEM, *p<0.05, **p<0.01, ***p<0.001,****p<0.0001, Student T-tests.

We can ascertain that the metabolic phenotypes observed in Th^Cre^; Npy^flox/flox^ mice are not caused by NPY^+^ neurons in the brain because firstly, no catecholaminergic neurons co-express NPY in hypothalamus^40^ or ventral tegmental area (VTA) (Supplementary Figure 14A-D), and *Npy* expression is not changed in the hypothalamus of Th^Cre^; Npy^flox/flox^ mice (Supplementary Figure 14E); secondly, even though there is catecholaminergic neurons in the hindbrain^41^, ablating NPY from the hindbrain has no effect on thermogenesis or BAT (Supplementary Figure 15A-G). The metabolic phenotype is also not caused by systemic NPY since NPY concentration in the blood plasma of Th^Cre^; Npy^flox/flox^ mice is unchanged (Supplementary Figure 14F). Altogether, we demonstrate that sympathetic-neuron derived NPY in adipose tissue is required for sustaining energy expenditure and leanness.

## Discussion

Mural cells are contractile cells that wrap the vasculature to regulate the diameter of vessels^42^ and have been described to be a source of thermogenic adipocytes both in WAT and BAT^3–5^, which are subject to the influence of NE released from adjacent sympathetic nerve terminals. These autonomic nerves co-release NPY^1,2,8^, but its function in the development and activity of thermogenic adipocytes was insofar unknown. In humans, mutations in NPY have been linked to high BMI but not an unhealthy dietary pattern^16^ (https://hugeamp.org/gene.html?gene=NPY, Supplementary Figure 16A). In mice, the loss of NPY receptors leads to late-onset obesity without increases in food intake^13–15^. Taken together, the experimental evidence points to an essential role of NPY in maintaining body weight homeostasis independent of regulating appetite. Three studies have implicated NPY in directing lipogenic actions in white adipocytes, which favoured the growth of white adipose mass^24,25,27^. However, some of these studies are not replicated here^24,25^, and others are not supported by modern evidence provided by our imaging data and several single-cell sequencing datasets^27^. Specifically, we show in our supplementary data that NPY neither directly promotes adipogenesis^24^ nor the proliferation of white adipocyte precursor cells^25^. Thus, the evidence disfavours a pro-obesity role for peripheral NPY, contradicting the current perception. We provide the first anatomical maps of NPY^+^ innervation in white and brown adipose tissues to conclude that only one-third of sympathetic innervation is neuropeptidergic and that it preferentially targets the arterioles where mural cells exist. Consistent with recent studies showing that sympathetic axons within adipose tissues retract and degenerate with the onset of obesity^7,43^, our data show that the local axonal source of NPY within these tissues is also obliterated, and this, again, refutes a direct adipogenic role of peripheral NPY. Moreover, several modern datasets omit NPY receptors in adipocytes or immune cells, whereas *Npy1r* expression is robustly detected in mural cells, identifying these cells as the postsynaptic targets of NPY^+^ innervation within adipose tissues. We show that NPY sustains the mural pool of the progenitor cells of thermogenic adipocytes by promoting their proliferation and facilitates the differentiation to thermogenic adipocytes *in vitro*. In vivo, NPY sourced from sympathetic neurons is necessary for supporting mural cells and the normal development of thermogenic adipocytes. Conditional knockout of NPY in sympathetic neurons (NPY-cKO) depletes mural cells and whitens brown fat before the onset of obesity, rendering this tissue hypertrophic and with reduced thermogenic ability. These NPY-cKO mice have lower energy expenditure, develop late-onset obesity on a regular chow diet and quickly develop obesity when fed a high-fat diet, although they do not eat more. This metabolic phenotype matches that of NPY receptors-whole-body-knockout mice^13–15^, which also develop late-onset obesity on a regular chow diet even though they eat less. Our results are the first to show the role of NPY in thermogenic adipocyte biology and clarify that central and peripheral sources of NPY have antipode roles in body weight homeostasis and the biological mechanism described in this study is a possible explanation for why numerous mutations of the appetite-stimulating NPY paradoxically associates with high BMI in humans (https://hugeamp.org/gene.html?gene=NPY)^16^ and highlights that energy dissipation may be more important than appetite in some individuals.

## Method

### 1. Animals

Th^Cre^ mice (B6.Cg-*7630403G23Rik^Tg(Th-cre)1Tmd^*/J; stock no. 008601), *Cx3cr1*^GFP/+^ mice (*Cx3cr1^tm1Litt^*/LittJ; stock no. 008451) were purchased from the Jackson Lab (JAX). Npy^flox/flox^ mice were a donation from Ivo Kalajzic Lab at the Department of Reconstructive Science, University of Connecticut^44^ under MTA. Tissues of NPY-GFP mice (B6.FVB-Tg(Npy-hrGFP)1Lowl/J) are from Tamas Horvath Lab at Brandy Memorial Laboratory, Yale University. Tissue of Npy1r^Cre^; Rosa26^tdTomato^ mice were from Professor Michael Roberts at Department of Otolaryngology-Head and Neck Surgery, University of Michigan. Sympathetic neuron-specific NPY-cKO mice were generated by crossing Th^Cre^ mice with Npy^flox/flox^ mice. Diet-induced obesity (DIO) was achieved by feeding mice an HFD (Diet Research, D12492) when they were 7 weeks old, and lasted for 10 weeks. The body weight of each mouse and the food consumption in each cage were recorded weekly. All the mice were group housed in standard housing under a 12-12-hour light-dark cycle and given access to diet and water *ad libitum*. All experimental procedures were performed on living animals in accordance with the United Kingdom ANIMALS ACTS 1986 under the project license (PPL number: P80EDA9F7) and personal licenses granted by the United Kingdom Home Office. Ethical Approval was provided by the Ethical Review Panel at the University of Oxford.

### 2. Energy expenditure and respiratory exchange rate measurement

Animals with 6 weeks were analyzed for Oxygen Consumption (V_O2_), Carbon Dioxide Production (V_CO2_), Energy Expenditure (EE) and Respiratory Exchange Rate (RER) using an indirect calorimetry system (Panlab; Harvard Apparatus; LE405 Gas Analyzer and Air Supply & Switching). Animals were maintained in individual cages, following a 12h light/dark cycle, with water and food supplied *ad libitum*, and controlled room temperature (21-23°C) and humidity. Metabolic data was collected for five days, after two-days acclimatation period. Animal’s body weight (g) and food (g) were measured before entering, and after exiting the cage. Results were normalized by weight, and graphs and statistical analysis were obtained using the CalR Web-based Analysis Tool for Indirect Calorimetry Experiments^45^.

### 3. Locomotor activity and body composition measurement

Animals with 8-9 weeks were analyzed for spontaneous locomotor activity (Panlab; Harvard Apparatus; LE001 PH Multitake Cage). Animals were maintained in individual cages, following a 12h light/dark cycle, with water and food supplied *ad libitum*, and controlled room temperature (21-23°C) and humidity. Data was collected for 72 hours, after one-day acclimatation period. Activity was recorded by COMPULSE v1.0 Software (PanLab). Animals with 7-9 weeks were analyzed for body composition using Minispec LF50 (Brucker).

### 4. Fasting and thermo-imaging

Animals with 12-14 weeks were used for thermos-imaging. Data in Figure 5E were recorded using Optris thermo-cameras (Optris PI 160 with standard 61° lens, Optris GmbH, Berlin, Germany), and Data in Supplementary Figure 15F were recorded using an FOTRIC 225s infrared camera and analysed using FOTRIC software. Animals were single-housed and put under the thermo-cameras, and their backs were shaved to expose skin above BATs. Animals were acclimatised for 4 days in individual cages with *ad libitum* access to food and water at room temperature under a 12/12 light-dark cycle before a 14-hour fast. Mice have *ad libitum* access to water during fasting.

BAT temperature was recorded every second for a 6-day period by storing the temperature of the warmest pixel in view using the software provided by the camera’s manufacturer (Optris PIX Connect, Optris GmbH). The BAT temperature at 0 and 14 hours was an average of temperatures during a1-hour period.

### 5. Stereotaxic injection

For viral injection, mice were anaesthetized with avertin (240mg/kg, i.p.) and fixed on a stereotaxic holder (RWD, Life Science, China). AAV2/9-hSyn-Cre-EGFP-WPRE-pA (Taitool, S0230-9, 2×1012 V.G./mL, 200nl for each side) or AAV2/9-hSyn-EGFP-WPRE-pA (Cat#S0237-9, Taitool, 2×1012 V.G./mL, 200nl for each side) was bilaterally injected into the NTS (NTS coordinate AP/ML/DV: –7.5/±0.35/-4.75 mm) of Npy^flox/flox^ mice.

### 6. Immunofluorescent staining (IF)

Superior cervical ganglia (SCG) and sympathetic axon bundles were dissected adipose tissues of mice perfused with 20 mL PBS and fixed in 4% paraformaldehyde (ThermoFisher, 043368.9M) for at least 4 hours. Cells collected after *in vitro* experiments were washed once with PBS and fixed using 4% PFA for at least 4 hours. Axon bundles were stained by first incubating with primary antibodies dissolved in the permeabilising buffer (3% BSA, 2% goat serum, 0.1% triton x-100, and 0.1% sodium azide in PBS) overnight at 4 □, followed by incubation with secondary antibodies and DAPI dissolved in the permeabilising buffer for 1 hour with gentle agitation. As for IF in fixed cells, cells were first incubated with the permeabilising buffer for 1 hour at room temperature (RT), and then stained at 4 □ overnight with primary antibodies diluted in the permeabilising buffer. Then, samples were washed and stained with DAPI and secondary antibodies for 1 hour at RT. Sympathetic fibres or fixed cells were then mounted on slides with Fluoromount-G mounting medium (ThermoFisher, 00-4958-02).

For cryosection, samples were first embedded in OCT (VWR, 361603E) and placed at –20□ until cryosectioning. Samples were cut into 10 µm-thick sections and thawed on charged SuperFrost Plus slides (VWR, 631-0108). Afterwards, the samples were stained using the same procedure as fixed cells. For whole-mount ganglia staining, PFA-fixed ganglia were dehydrated in 30%, 50%, 70%, and 90% once and twice in 100% ethanol, and the ganglia were kept shaking for 20 minutes at each concentration, and then ganglia were rehydrated in 90%, 70%, 50%, and 30% ethanol. Afterwards, ganglia were digested with 1.5 U/mL dispase-1 (Roche-04942078001), 0.5 mg/mL collagenase (Sigma, C2674), and 300 μg/mL hyaluronidase (Sigma, H3884) diluted in PBS, shaking in 37 □ water bath for 30 minutes. Ethanol-treated ganglia were blocked in blocking solution (3% BSA, 2% goat serum, 0.1% triton x-100, and 0.1% sodium azide in PBS) for 2 hours and then incubated with primary antibodies diluted in the permeabilising buffer for 3 days at 4□. 3 days later, ganglia were incubated in secondary antibodies diluted in the permeabilising buffer for another 3 days at 4□. Finally, ganglia were dehydrated in 30%, 50%, 70%, 90% ethanol and twice in 100% ethanol, and were cleared using ethyl cinnamate (ECi). Cleared ganglia were mounted in a coverslip-slide sandwich filled with ECi.

Images were acquired using a Zeiss LSM880 confocal microscope with Zen-black software (V2.1). The samples were imaged using a 10x/0.45NA objective, with a 1.04 μm, 1.04 μm, 3.64 μm xyz voxel size, a 20x/0.8NA objective, with a 0.52 μm, 0.52 μm, 1.00 μm xyz voxel size, or a 63x/1.4NA oil-immersion objective, with a 0.13 μm, 0.13 μm, 1.00 μm xyz voxel size. Solid-state 405nm, 561nm, 633nm, and Argon 488 lasers were used for DAPI, AF488, AF546, and AF647 fluorophores respectively.

### 7. Adipose Tissue Clearing

Whole fat pads were stained and cleared using a modified version of iDISCO as previously described^18,46^. Briefly, white adipose tissues (WATs) were dissected from mice perfused with 20 mL PBS and fixed with 4% paraformaldehyde overnight. Tissues were pre-treated with 20%, 40%, 60%, 80% and twice in 100% methanol for 30 minutes each at RT, and then bleached using 5% H_2_O_2_ diluted in 100% methanol for 24 hours at 4 □ shaking. Bleached samples were rehydrated using 80%, 60%, 40%, and 20% methanol for 30 minutes each and then in PTwH buffer (0.2% Tween-20, 10 μg/mL heparin, and 0.02% sodium azide in PBS) for 30 minutes. Afterwards, samples were incubated in permeabilising solution (20% DMSO, 0.2% Triton x-100, 0.3M glycine in PBS) at 37 □ water bath overnight and blocked using blocking buffer (10% DMSO, 2% Triton x-100, 0.02% sodium azide and 5% goat serum in PBS). For immunolabeling, samples were incubated in primary antibodies diluted in antibody dilution buffer (5% DMSO, 5% goat serum, 0.2% Tween-20 and 10 μg/mL heparin in PBS) at 37□ shaking for 5 days, then washed with PTwH for 1 day, and then incubated with secondary antibodies and DAPI diluted using antibody dilution buffer at 37□ shaking for another 3 days. Immunolabelled tissues were embedded in 1% agarose and dehydrated in 20%, 40%, 60%, 80%, and 100% methanol for 1 hour each, and then 100% overnight at room temperature. Samples were incubated in dichloromethane (DCM) (Sigma, 270997) until they sank, and then incubated in dibenzyl ether (DBE) (Sigma, 179272) until clear. Samples were stored in DBE at room temperature, and transferred into ECi before imaging. Images of cleared tissues were required using a Myltenyi Biotec Ultramicroscope II light-sheet microscope. The step size was 8μm; the thickness of the light sheet was 3.98 μm; the horizontal dynamic focusing was set to 8 steps; and the exposure time was set at 180ms. The samples were illuminated using a bi-directional light-sheet and scanned under a 2x/0.5 NA objective, with a 4.03 μm, 4.03 μm, 8 μm xyz voxel size. A 488nm laser with a 525/50 filter, a 561nm laser with a 620/60 filter, and a 638nm laser with a 680/30 filter were used for AF488, AF546, and AF647 respectively.

### 8. Antibodies

The following primary antibodies were used for immunofluorescent staining: rat anti-CD31 (BioLegend, 102501, MEC13.3,1:100 dilution), rat anti-PLVAP (BioLegend, 120503, MECA32, 1:100 dilution), rabbit anti-DES (Abcam, AB15200, 1:500 dilution), rabbit anti-NPY (Cell Signalling, D7Y5A, 1:500 dilution), rabbit anti-NPY (Abcam, AB30914, 1:500 dilution), chicken anti-TH (Aves Labs, TYH73787982, 1:500 dilution), rabbit anti-TH (Sigma, Ab152, 1:500 dilution), mouse anti-NPY1R (Santa Cruz, sc-393192, 1:200 dilution), rat anti-PDGFRα (BioLegend, 135902, APA5, 1:200 dilution), rabbit anti-TAGLN (Abcam, AB14106, 1:250 dilution), Cy3 anti-αSMA (Sigma, C6198, 1A4, 1:250 dilution), goat anti-SOX17 (R&D, AF1924, 1:250), goat anti-EPHB4 (R&D, AF3034, 1:250).

The following antibodies were used for FACS and flow cytometry: AF700 anti-CD45 (BioLegend, 103128), BUV395 anti-CD45 (BioLegend, 564279), Pacific Blue anti-CD31 (102421), APC anti-PDGFRa (BioLegend 135907), AF488 anti-NG2 (Sigma, MAB5384A4), and AF488 anti-DES (Abcam, AB185033, Y66). LIVE/DEAD™ Fixable Near-IR Dead Cell Stain Kit (ThermoFisher, L10119) was used for live/dead staining.

### 9. qPCR

RNA extraction was performed using Trizol reagent (Thermofisher, 15596026), cDNA was synthesized using SuperScript II Reverse Transcriptase (Invitrogen, 18064022), and qPCR was done using Power SYBR Green PCR Master Mix (LifeTech 4368706).

The following primers were used in this research:

**Table 1:**
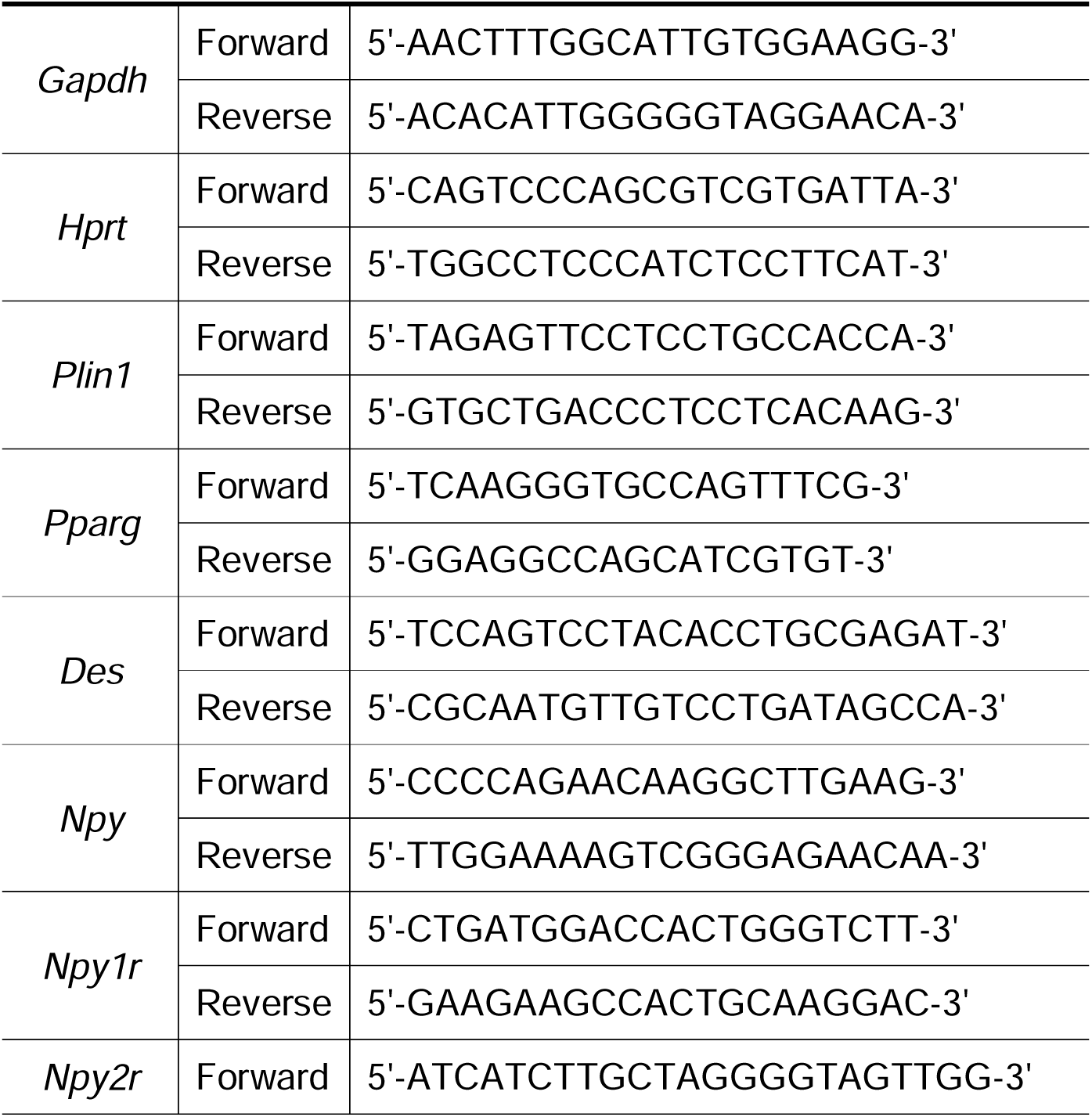
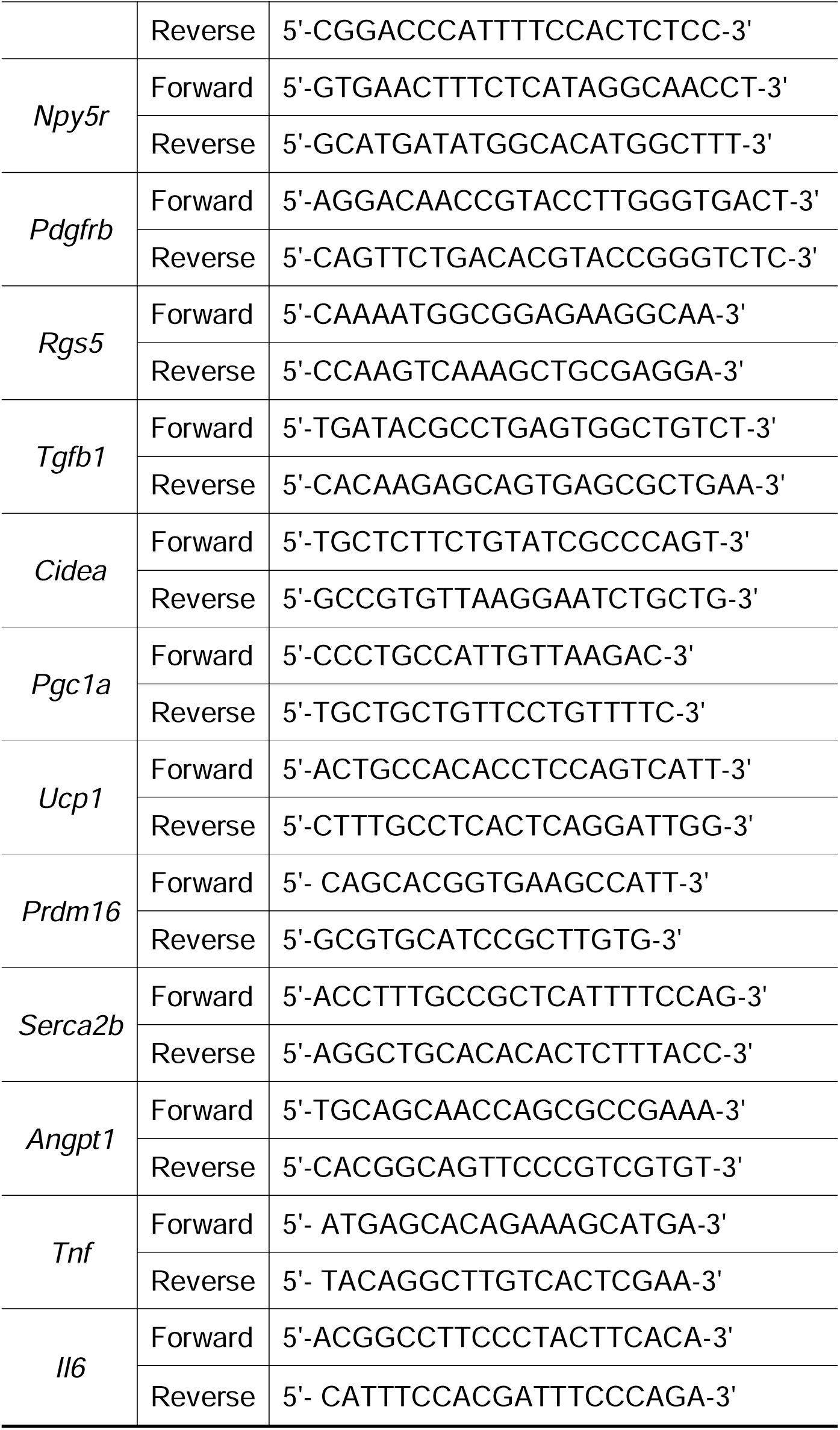
The Sequence of Primers for RT-qPCR.

ΔCt method was used to quantify the gene expression using the formula: relative expression = 2^(-(Ct_target gene_-Ct_reference gene_)).

qPCR data shown in Supplementary Figure 15 was collected using CFX96 Real-Time PCR Detection System (Bio-Rad) with TB Green Premix Ex Taq II (Cat# RR820A, TaKaRa), and ΔΔCt was used for calculating fold changes in gene expression to compare Hindbrain: Npy^flox/flox^; AAV-CRE and WT mice.

### 10. Protein extraction and blood plasma isolation for NPY ELISA

Proteins were extracted from iWATs as previously described in a published paper^47^. Briefly, dissect iWATs were put in homogenising tubes filled with 300 μL/50mg RIPA lysis buffer (Sigma, 20188) containing 50 μM DPP-IV inhibitor (Sigma, DPP4-010) and 500 KIU/mL Aprotinin (Sigma, A6103-1MG) and homogenised using Precellys 24 homogeniser. The lipid in the tissue homogenise was removed by centrifuging the tissue homogenise at 20000x rcf at 4□ for 15 minutes, and the clear part was kept. The process was repeated 3 times to completely remove lipids.

To do terminal blood collection, the mice were euthanised by i.p. injection with 10 μL/g pentobarbital, and then blood was collected from the left ventricle using 25G needles and syringes coated with 100 mM EDTA. The blood was then put into tubes with 5 μL 100 mM EDTA and centrifuged at 1000x rgf at 4□ for 15 minutes to separate blood plasma from blood cells, and then the supernatant was moved to new tubes with DPP-IV inhibitor (final concentration 50 μM) and Aprotinin (final concentration 500 KIU/mL). The concentration of NPY in the iWAT and blood plasma of mice was determined using an NPY ELISA kit (Merck, EZRMNPY-27K).

### 11. Single-cell suspension and flow cytometry

Dissected adipose tissues were minced and digested in the enzyme mixture (for each sample, 500μL Collagenase II (4 mg/mL, C6885), 500μL Hyaluronidase (5.3 mg/mL=40000 U/mL, H3884), and 5 μL Dnase I (BioLabs, M0303L)) in a 37 □-water bath shaking for 45 minutes, and samples were pipetted every 10 minutes. Digestion was stopped by adding FACS buffer (PBS containing 2% FBS), and single-cell suspension was collected by filtering the digested sample using EASYStrainer cell sieves with 70μm mesh (Greiner, 542070).

To prepare the samples for sorting or flow cytometry, cells were first treated with red-blood-cell lysis buffer (BioLegend, 420301) to remove red-blood cells, and treated with Fc block (ThermoFisher, 14-9161-73) before staining with antibodies. Before immunolabelling for intracellular markers, cells were fixed and permeabilised using the eBioscience Intracellular Fixation and Permeabilization Buffer Set (ThermoFisher, 88-8824-00). Cell sorting was done using a BD FACSAria III sorter, and flow cytometry data were acquired using a BD FACSAria III sorter or a BD LSRFortessa X20 cytometer with a BD FACSDiva v6.0 software. Cytometry data were analysed using FlowJo v10.8.1.

### 12. Isolation and primary culture of mural cells and *in vitro* stimulation

Mural cells were isolated and cultured following a modified version of the protocol previously published^48^. Briefly, the mice were euthanised by i.p. injection with 10 μL/g pentobarbital and perfused with 15mL PBS to remove blood. The stromal vascular fraction (SVF) of the BATs was isolated using the method described above. Then, the cells were seeded in 10 cm dishes with 10 mL DMEM with 10% FBS for 2 days for the cells to attach to the bottom. Afterwards, 5mL of the medium was replaced with mural cell-specific medium (DMEM with 2% FBS supplied with 1% mural cell-growth supplement (Science Cell, 1252)). 5 mL of the medium in the dishes was refreshed every 2 days until the cell grow confluent. To get pure mural cells, cells were labelled with an APC anti-CD104b antibody (BioLegend, 136007), and mural cells were sorted using MojoSort^TM^ magnetic sorting protocol with MojoSort^TM^ buffer (BioLegend, 480017) and anti-APC nanobeads (BioLegend, 480071). To perform *in vitro* co-culturing experiments. The cells were seeded at 5×10å4 cells/mL on glass coverslips and indicated concentration of PDGF-BB (R&B, 220-BB) and NPY (Cayman Chemical, CAY15071) were added to the cells 24 hours after the cells were seeded. Finally, the cells were collected 5 days later and either fixed with 4% PFA for imaging or lysed with Trizol for RNA extraction and qPCR.

### 13. Cell lines

RAW264.7 murine macrophage cell line (Sigma, 91062702) was purchased from Sigma. The complete medium for culturing the cells includes high glucose DMEM with glutamate (Sigma, 41965039) and 10% FBS (Sigma, 12133C).

The 3T3-L1 preadipocyte cell line was a gift from Associate Professor Robin Klemm at the Department of Physiology, Anatomy and Genetics, University of Oxford. The complete medium for culturing the cells includes high glucose DMEM with glutamate (ThermoFisher, 41965039) and 10% calf serum (Sigma, 12133C). To subculture preadipocytes, the medium was moved, and cells were washed with 1mL pre-warmed 0.05% trypsin (ThermoFisher, 25300054), and digested with 200μL trypsin at 37 □ for 5-10 minutes. Afterwards, digestion was stopped, and cells were resuspended using the complete medium.

### 14. Induction to white and thermogenic adipocytes

To differentiate 3T3-L1 cells to white adipocytes, cells were seeded in 12-well plates at a density of 1×10å5 cells/mL. Medium was refreshed until the cells were confluent. 2 days after the cells were confluent, induction medium (10% FBS, 500μM 3-Isobutyl-methylxanthine and 1 μM dexamethasone in DMEM) with or without 1 μg/mL insulin were added into each well. 3 days later, the induction medium was replaced with the maintenance medium (DMEM with 10% FBS and with or without 1 μg/mL insulin). Afterwards, medium was refreshed with the maintenance medium every 2 days. The total differentiation time for each well was 8 days. NPY treatments in the experimental groups starts when the induction medium was added and lasted through the whole differentiation process.

SVF and mural cells was differentiated to thermogenic adipocytes as previously published^5,30^. SVF and mural cells was isolated and seeded in 12-well plates at a density of 1×10å5 cells/mL. When cells reach 95% confluency, beige cell induction medium (DMEM with 10% FBS, 125 μM Indomethacin, 5 μM Dexamethasone, 500 μM 3-Isobutyl-methylxanthine, and 0.5μM Rosiglitazone) was added to each well. 2 days later, the induction medium was replaced with maintenance medium (DMEM with 10% FBS, 5 μg/mL Insulin, and 1 nM Triiodothyronine). Afterwards, the maintenance medium was refreshed every 2 days. The total differentiation time for each well was 8 days. NPY treatments in the experimental groups started when the induction medium was added and lasted through the whole differentiation process. Oxygen consumption was measured using a Seahorse XF Analyzer.

### 15. Cell proliferation assay

Cell proliferation assay was done with Click-iT EdU Cell Proliferation Kit for imaging (C10340). Mural cells were seeded at 0.5×10å5 cells/ml in 12-well plates on coverslips and cultured overnight before the experiments. Afterwards, cells were labelled with EdU and cultured with or without 1 μM NPY for 6 hours in medium for mural cells (low-glucose DMEM with 2%FBS). Cells were then fixed with 4% PFA and permeabilising buffer for 30 minutes respectively, and EdU was labelled with AF647 using Click-iT® Plus reaction cocktail. After IF staining with anti-DES, cells were imaged using a Zeiss LSM880 confocal microscope with Zen-black software (V2.1), with a 20x/0.8NA objective, a 0.52 μm, 0.52 μm, 1.00 μm xyz voxel size.

### 16. scRNA-Seq dataset analysis

Public scRNA-Seq datasets were downloaded from GEO, and were analysed using the following method. Cells with <200 unique detected genes or >5% mitochondrial counts were discarded. After filtering, the gene x cell matrix was normalised using ‘NormalizeData()’ in Seurat v4.2.0^49^ in R (v4.2.2) environment. Then, the data were scaled using ‘ScaleData()’, and we performed linear dimensional reduction by PCA and calculated UCMP coordinates for all cells using Seurat v4.2.0. The cells were clustered using ‘FindNeighbour()’, with dimensions set to 15 and ‘FindClusters()’, with resolution set to 0.5. Each cluster were identified based on differentially expressed genes and known markers in published literatures^4,22,29,50–52^.

### 17. Figure quantification

The overlapping between TH, NPY and CD31 was calculated using JACoP^53^, a plugin in Fiji^54^. To make this calculation, 472.33 μm, 472.33 μm, 30 μm xyz regions were randomly picked and maximally project to z using Fiji script. The labelled areas were automatically segmented using threshold set by default, and the overlapping percentages were calculated automatically by JACoP. 5 views from 3 biological replicates for iWAT were quantified.

The innervation of NPY^+^ axons was quantified using the “Surface” tool in ImarisV9.2. The labelled area of NPY^+^ axons and CD31^+^ vessels in whole cleared iWATs were automatically segmented, and the innervation of NPY in vasculature was calculated as Volume_NPY_^+^/Volume_CD31_^+^. The coverage of DES^+^ mural cells in iWAT was calculated similarly as Volume ^+^/Volume ^+^. Confocal images showing NPY^+^ innervation were quantified using Fiji with an automatic unbiased method. Area_NPY+_ and Area_CD31+_ were segmented using Otsu thresholding method and measured using “measure” program in Fiji.

The percentage of EdU^+^ cells were counted with an unbiased automatic method using “detect particles” in Fiji^54^. Threshold was set automatically using Otsu thresholding method. 6 views from 6 biological replicates were used for quantification.

The percentage of PDGFRα^+^ and DES^+^ cells were count manually based on the PDGFRα and DES signals. The percentage of TH^+^ and NPY^+^ neurons in hypothalamus and VTA was also counted manually. To calculate the ganglionic mural cell coverage, 531 μm, 531 μm, 30 μm xyz views were randomly picked and maximally project to z using Fiji script. The labelled areas were segmented using threshold set by default, and the coverage was calculated as Area_NPY_^+^/Area_CD31_^+^. 11 views from 4 biological replicates were quantified for each condition.

The size of adipocytes in WAT and BAT was quantified as previously described^55^ in Fiji. Briefly, WAT and BAT were processed to paraffin-embedded sections (3 μm sections), stained with H&E and scanned into digital images using a NanoZoomer-SQ Digital slide scanner –Hamamatsu. Images were randomly picked and the plugin Analyse Particles was used to count the number and measure the size of adipocytes and droplets in BAT. For each BAT, 11000 lipid droplets were measured.

### 18. Statistics

GraphPad Prism v.9.5.0 was used for statistical analysis. Mean and standard error of the mean (SEM) were used to represent a sample. Chi-Square Test and paired two-tailed Student T-test were used to compare two samples; ANOVA was used to compare multiple data points.

## Data Availability

The scRNA-Seq datasets of iWAT and BAT were deposited on the Gene-Expression Omnibus under the accession numbers GSE154047 and GSE160585. All the numeric data for supporting this research are available within the paper and the Supplementary information.

## Code Availability

The code used for scRNA-Seq analysis is deposited at Github (https://github.com/PLAVRVSO/scRNA-Seq-analysis-of-BAT-SVF).

## Acknowledgement

We thank I.Kalajzic for sharing the Npy^flox/flox^ mice; M.Silveira and M.Roberts for sharing the Npy1r^Cre^; Rosa26^tdTomato^ mice; T.Hovarth for sharing the NPY-GFP mice; R. Klemm for the 3T3-L1 cell line; V.Vyazovskiy and S.Wilcox for access to thermo-camera and technical assistance; M.Dustin at the Kennedy Institute of Rheumatology for access to light-sheet microscope, especially C.Lagerholm for his technical assistance; the Dunn School of Pathology for confocal microscopy, especially A.Wainman for his technical assistance; L. Zhou and X. Lu for their help with histology; K.Zhang for her technical assistance; all the members of Domingos Lab for discussions and advice on the project.

Y. Zhu receives scholarship from Shineroad Industry Developments Co. Ltd. L. Yao is supported by China Scholarship Council. J. Chen is supported by National Natural Science Foundation of China (No. 32100821). This research is funded by the Wellcome Trust –Howard Hughs Medical Institute International research scholar award (208576/Z/17/Z), the ERC consolidator award (ERC-2017 COG 771431), and the Pfizer ASPIRE Obesity award.

## Contributions

Y.Z designed and performed all experiments with guidance from A.D unless otherwise stated. Y.Z wrote the first draft of the manuscript and A.D. wrote the subsequent versions with Y.Z. L.Y assisted Y.Z and performed experiments shown in Supplementary Figure 2, 6, and 7G with guidance from Y.Z and A.D. A.L.G, B.B, and A.R.S conducted experiments shown in Figure 5D, Supplementary Figure 13A-E with guidance from L.V. G.S helped Y.Z conceive this project and provided advice on scRNA-Seq data analysis. A.C acquired light-sheet images at the Francis Crick Institute on samples cleared and stained by Y.Z shown in Supplementary Figure 9A&C and gave technical advice on light-sheet imaging. N.M. helped with the colony management. C.L imaged the first samples cleared and stained by Y.Z (not shown in this manuscript). M.D permitted access to the Kennedy Imaging Facility; K.A permitted access to the Crick Imaging Facility. T.H provided the brains of NPY-GFP mice shown in Supplementary Figure 14. S.K gave intellectual contributions pertaining to beige fat biology, and I.A. conducted the experiment shown in Figure 3C-G with guidance from S.K. J.C conducted the experiments shown in Supplementary Figure 15 with guidance from C.Z.

## Competing Interest Declaration

All authors declare no competing interest.

**Supplementary Figure 1.**
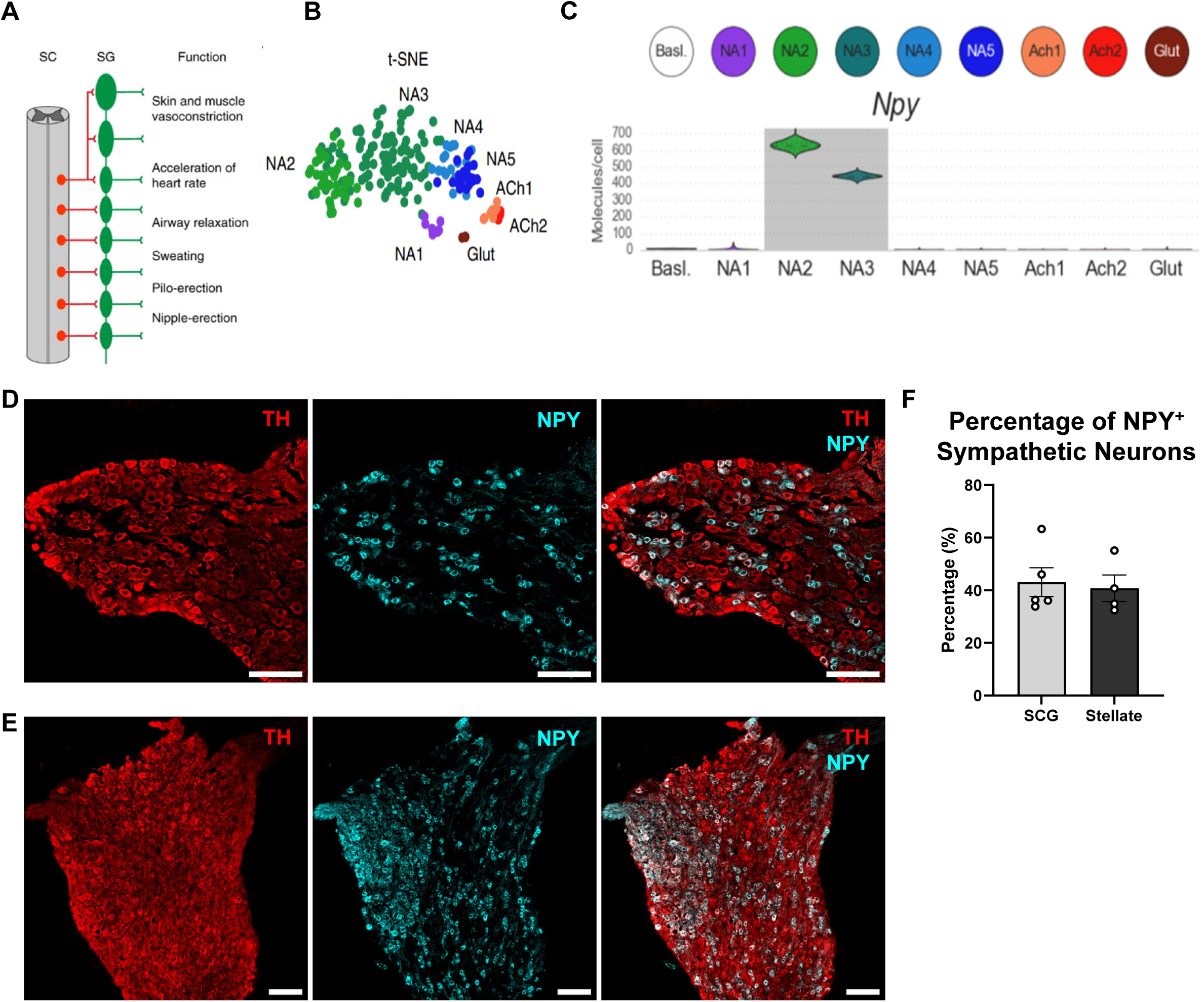
*Npy* is expressed by a subpopulation of sympathetic neurons. (A) Schematic showing the tissues innervated by each sympathetic ganglia^17^. (B) Single cell RNA-Seq dataset of sympathetic ganglia chain published by Linnarsson Lab to show the clustering of neurons^17^.(C) Violin plot showing the expression of *Npy*. (D-E) Confocal image of cryo-sectioned (D) superior-cervical ganglion (SCG) and (E) stellate ganglia (SG) of ND-treated WT male 8-week-old mice stained with anti-TH (red) and anti-NPY (cyan). Scale bars=100 μm. (F) The percentage of NPY+ sympathetic neurons quantified based on images as in D&E (n=5). All values are expressed as mean ± SEM.

**Supplementary Figure 2.**
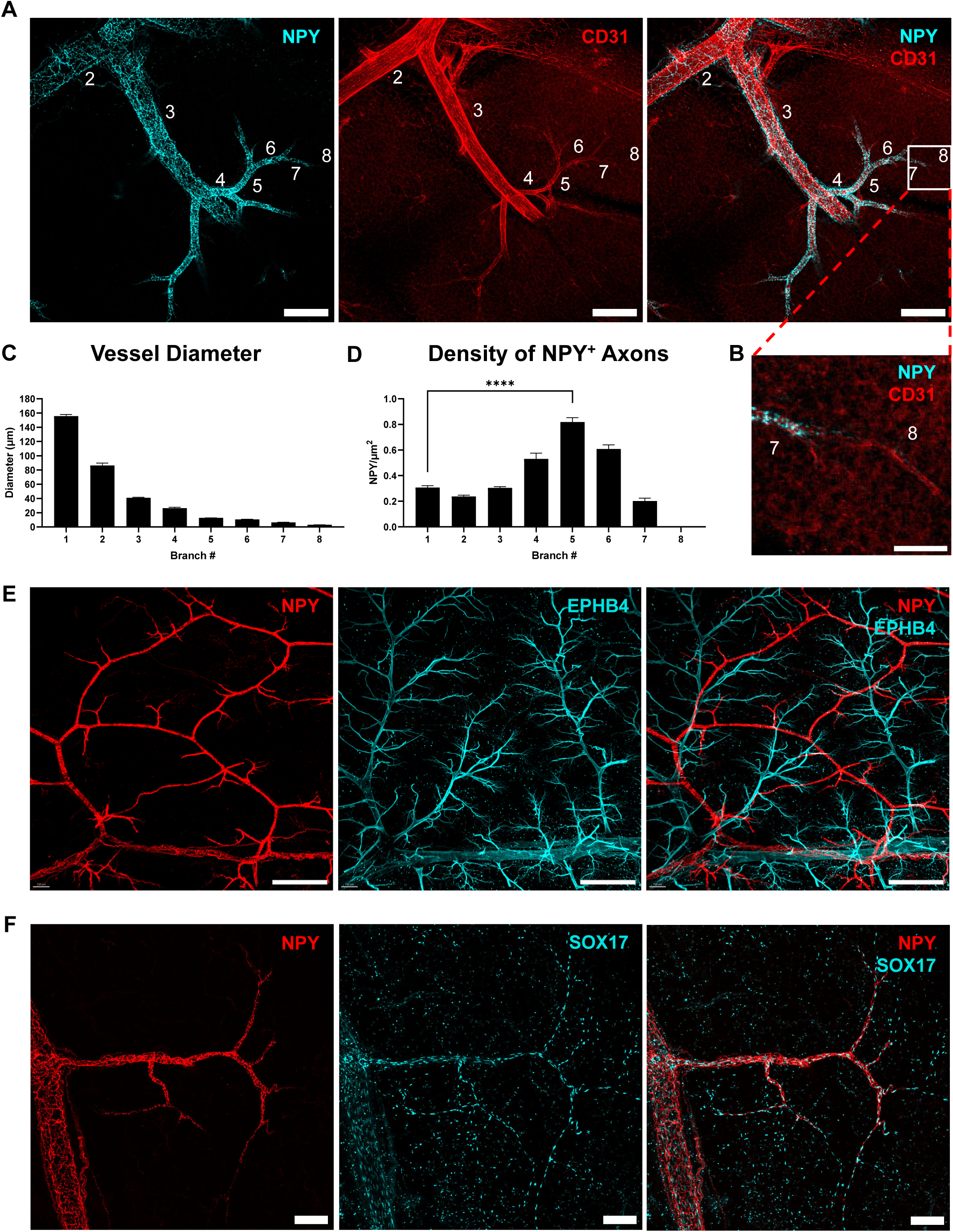
NPY^+^ axons preferentially innervate 5^th^ order of arteriole branches. (A) Confocal images of cleared adipose tissue stained with anti-NPY (cyan) and anti-CD31 (red). Number indicates the branch orders. Scale bar = 150 μm. (B) Zoom-in of (A), scale bar = 50 μm. (C) The diameters of vessels and (D) the density of NPY+ axons at each branch level quantified based on images as in A (n=6). (D) Light-sheet images of cleared adipose tissue stained with anti-NPY (red) and anti-EPHB4 (cyan). Scale bar = 500 μm. (F) Confocal images of cleared adipose tissue stained with anti-NPY (red) and anti-SOX17 (cyan). Scale bar = 100 μm. All values are expressed as mean ± SEM, *p<0.05, **p<0.01, ***p<0.001, ****p<0.0001, Student T-tests.

**Supplementary Figure 3.**
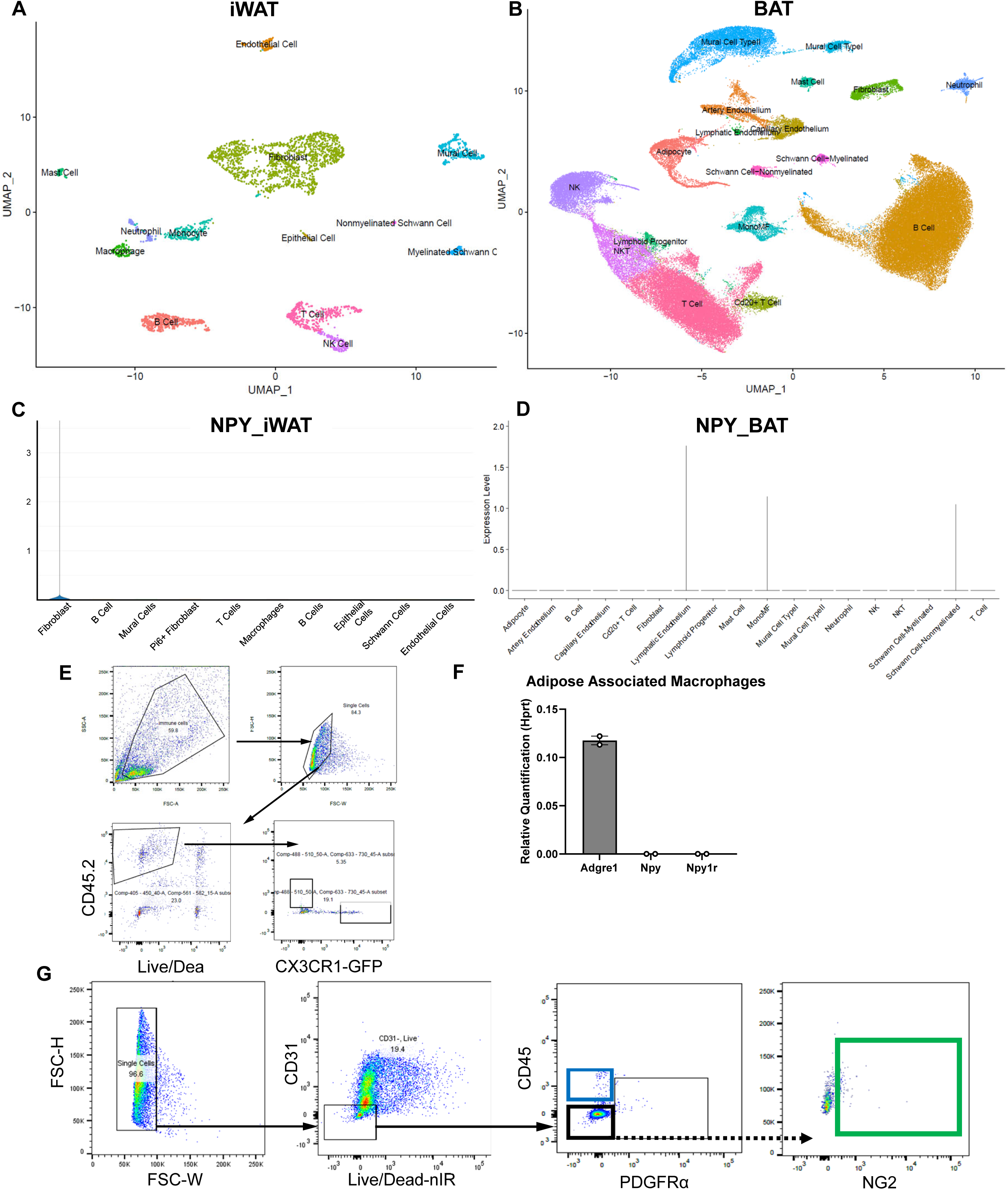
*Npy* is not expressed in adipose tissue, and NPY receptors are not expressed in macrophages. (A-B) Embedding plots showing different clusters of cells in the SVF of murine (A) iWAT^22^ and (B) BAT^5^. (C-D) Violin plot showing the expression of *Npy* in the SVF of murine (C) iWAT and (D) BAT. (E) Gating strategy of sorting adipose associated macrophages (ATM) from *Cx3cr1*^GFP/+^ reporter mice. ATMs were sorted as live, CD45.2^+^, GFP^+^ cells. (F) The expression of *Adgre1* (F4/80), *Npy*, and *Npy1r* in adipose associated macrophages (ATM) (n=2). (G) Gating strategy of sorting mural cells from the SVF of the iWAT of wild type normal-diet treated mice. mural cells (green box) are sorted as live, CD31^−^/CD45^−^/PDGFRα^−^/NG2^+^ cells; immune cells (blue box) are sorted as live, CD31^−^/CD45^+^ cells. All values are expressed as mean ± SEM, *p<0.05, **p<0.01, ***p<0.001, ****p<0.0001, Student T-tests.

**Supplementary Figure 4.**
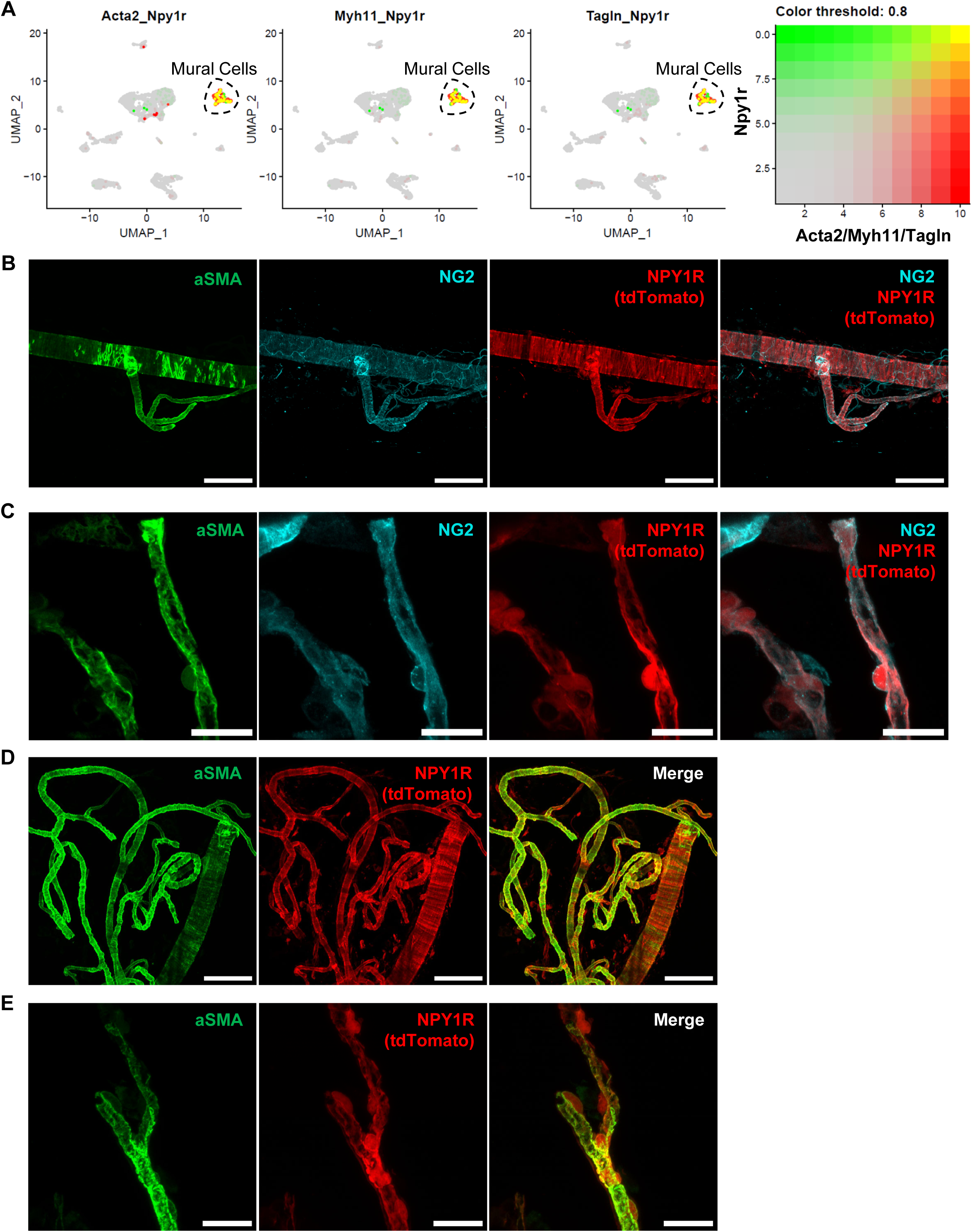
NPY1R-tdTomato is colocalised with NG2 and aSMA in mural cells. (A) Embedding plot showing the co-expression of mural markers *Acta2* (aSMA), *Myh11*, and *Tagln* (red) with *Npy1r* (green). Co-expression is in yellow^22^. (B) Confocal images of vessels dissected from adipose tissue of Npy1r^Cre^; Rosa26^tdTomato^ mice stained with anti-aSMA (green), scale bar=100 μm. (C) Zoomed-in images of vessels as in B, scale bar=20 μm. (D) Confocal images of vessels dissected from Npy1r^Cre^; Rosa26^tdTomato^ mice stained with anti-aSMA (green) and anti-NG2 (cyan), scale bar=100 μm. (E) Zoomed-in images of vessels as in D, scale bar=20 μm.

**Supplementary Figure 5.**
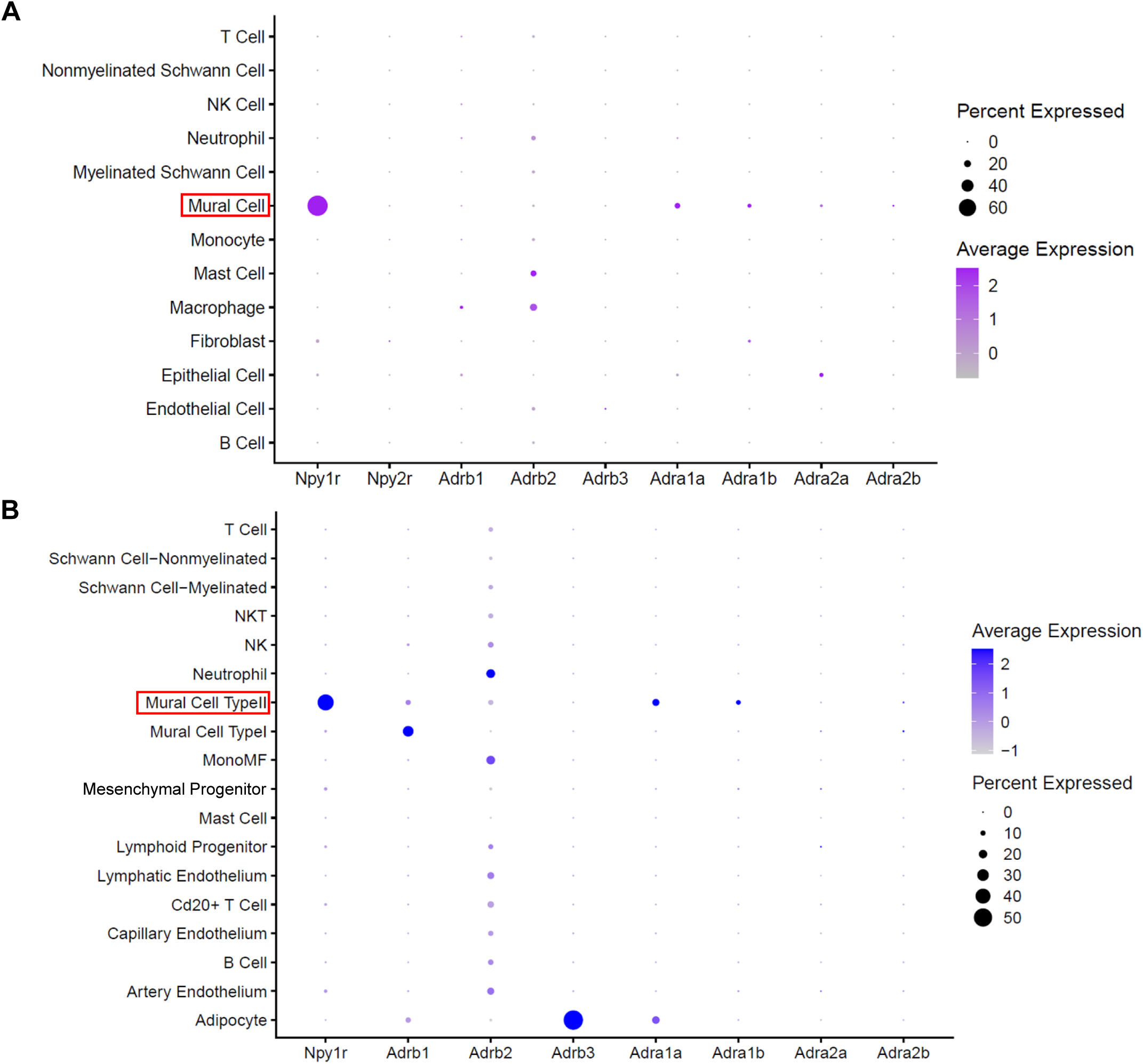
The expression levels of *Npy2r* and *Npy5r* in adipose tissue are negligible. (A) Dot plot showing the expression of *Npy1r*, *Npy2r*, *Adra1a*, *Adra1b*, *Adra2a*, *Adrb1*, *Adrb2*, and *Adrb3* in different cells in the SVF of mice iWAT. (B) Dot plot showing the expression of *Npy1r, Adrb1, Adrb2, Adrb3, Adra1a, Adra1b, Adra2a,* and *Adra2b* in different types of cells in the SVF of mouse BAT. The size of the dot represents the percentage of cells expressing a certain gene, and the darkness of the dot represents the expression level. Mural cell clusters are highlighted with red rectangle.

**Supplementary Figure 6.**
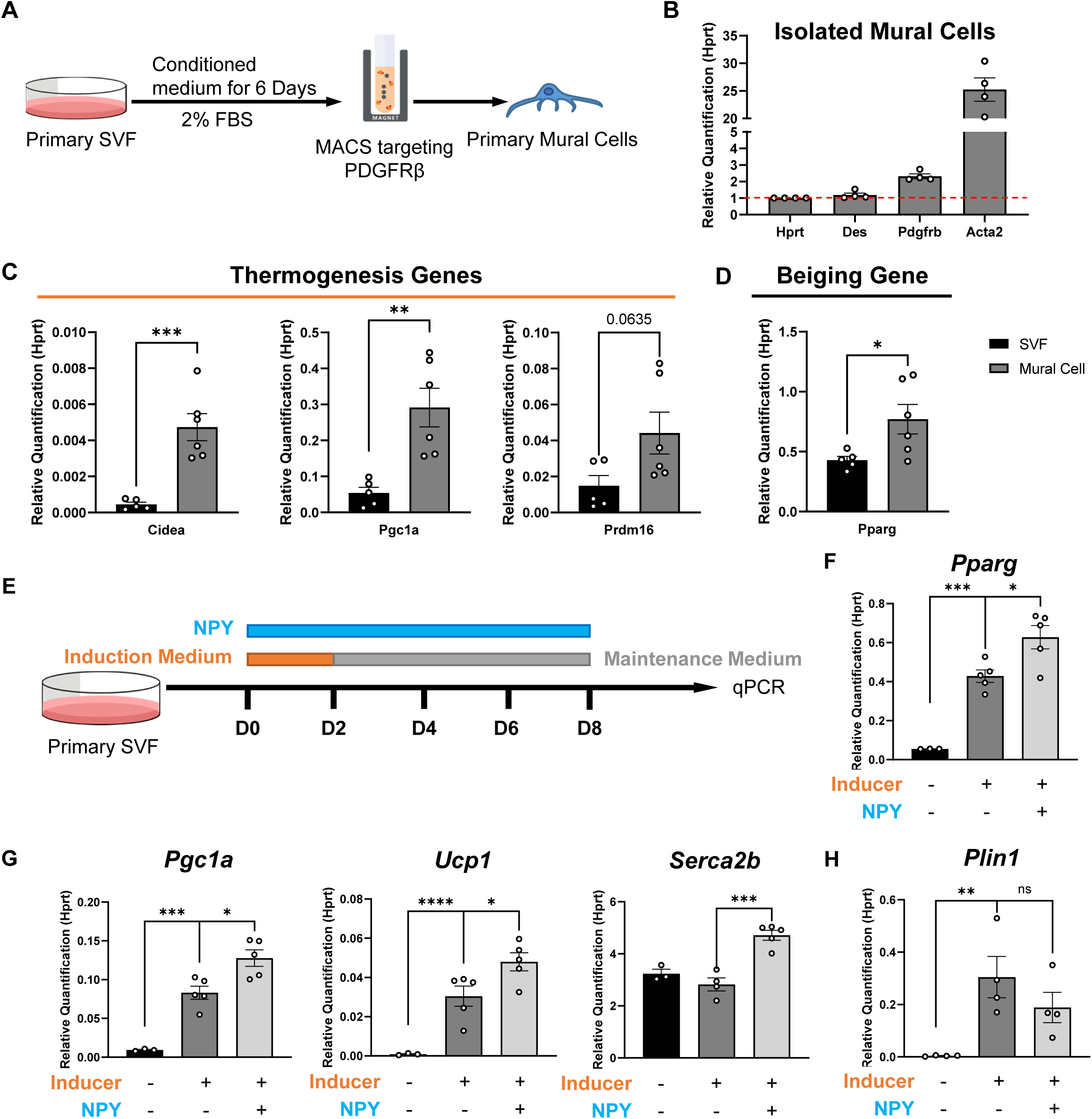
Mural cells can differentiate into thermogenic adipocytes, and NPY facilitates the differentiation of SVF to thermogenic adipocytes. (A) Schematic of isolating mural cells from adipose tissue. (B) The expression level of mural cell markers *Des*, *Pdgfrb*, and *Acta2*. Red dashed line indicates the expression level of reference gene *Hprt*. (C-D) The expression level of (C) thermogenesis genes *Cidea*, *Pgc1a*, and *Prdm16*, and (D) beiging gene *Pparg* in adipocytes differentiated from the stromal-vascular fraction (SVF) of adipose tissue and isolated mural cells. (E) Schematic showing *in vitro* method to differentiate primary stromal vascular fraction (SVF) of iWATs to thermogenic adipocytes. (F-H) The expression of (F) beiging gene *Pparg* (n=3&5&5), and (G) thermogenesis genes *Pgc1a, Ucp1, Serca2b* (n=3&5&5), and (H) adipogenesis gene *Plin1* (n=4), in thermogenic adipocytes differentiated from primary SVF. All values are expressed as mean ± SEM, *p<0.05, **p<0.01, ***p<0.001, ****p<0.0001, Student T-tests.

**Supplementary Figure 7.**
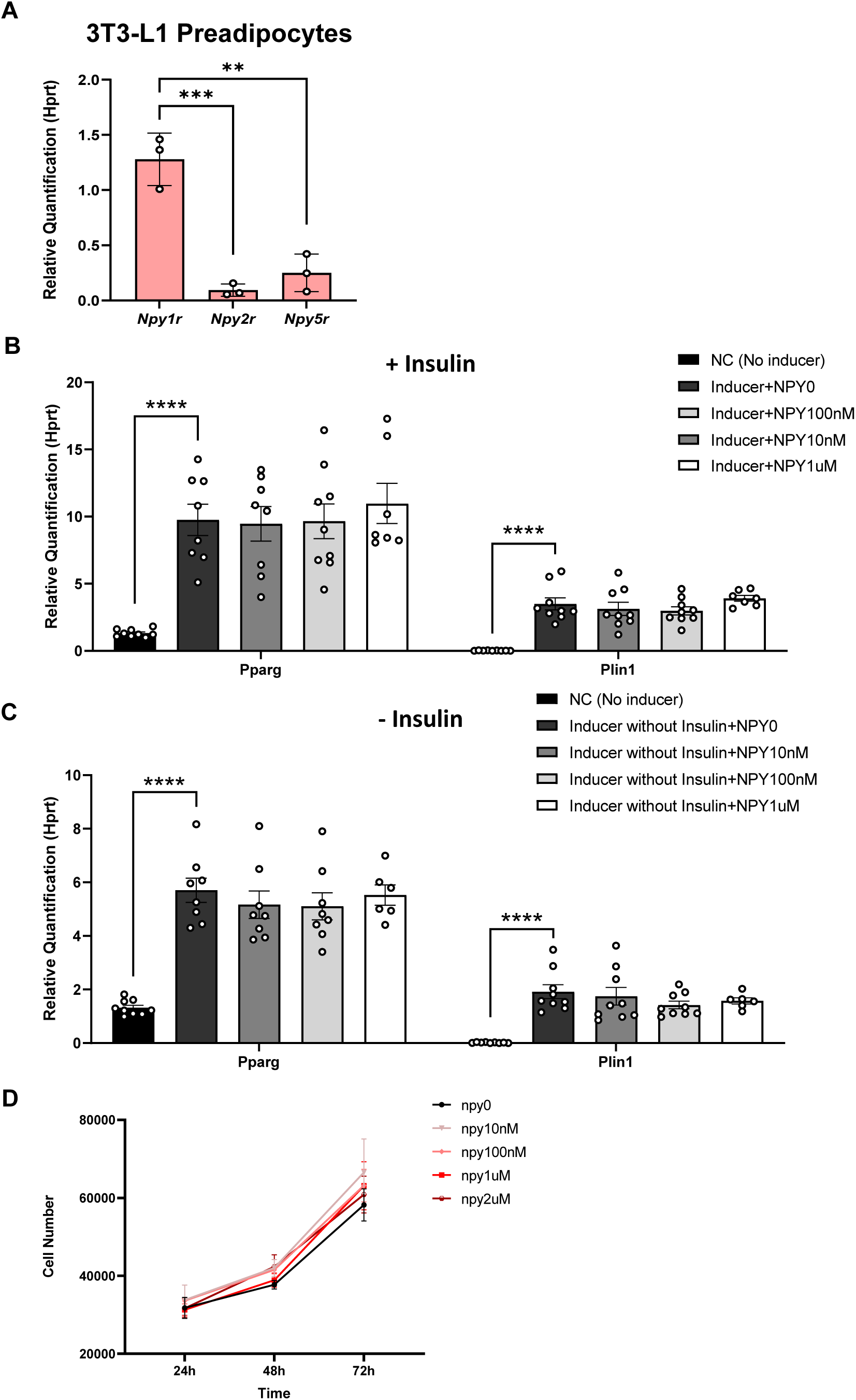
NPY does not affect general adipogenesis or the proliferation of 3T3-L1. (A) The expression of *Npy1r*, *Npy2r*, and *Npy5r* in 3T3-L1 preadipocyte cell line (n=3). (B-C) The expression of adipogenesis markers *Pparg* and *Plin1* in differentiated 3T3-L1 preadipocytes induced by induction medium (B) with or (C) without 1 μg/mL insulin (n=9, 8, 8, 9 & 7 for *Pparg*, n=9, 9, 9, 9 & 7 for *Plin1*). The concentration of NPY is indicated in the plot. (D) The impact of NPY on the proliferation of 3T3-L1 preadipocytes revealed by the cell number after incubating with or without NPY for 24, 48, and 72 hours (n=3). All values are expressed as mean ± SEM, *p<0.05, **p<0.01, ***p<0.001, ****p<0.0001, Student T-tests.

**Supplementary Figure 8.**
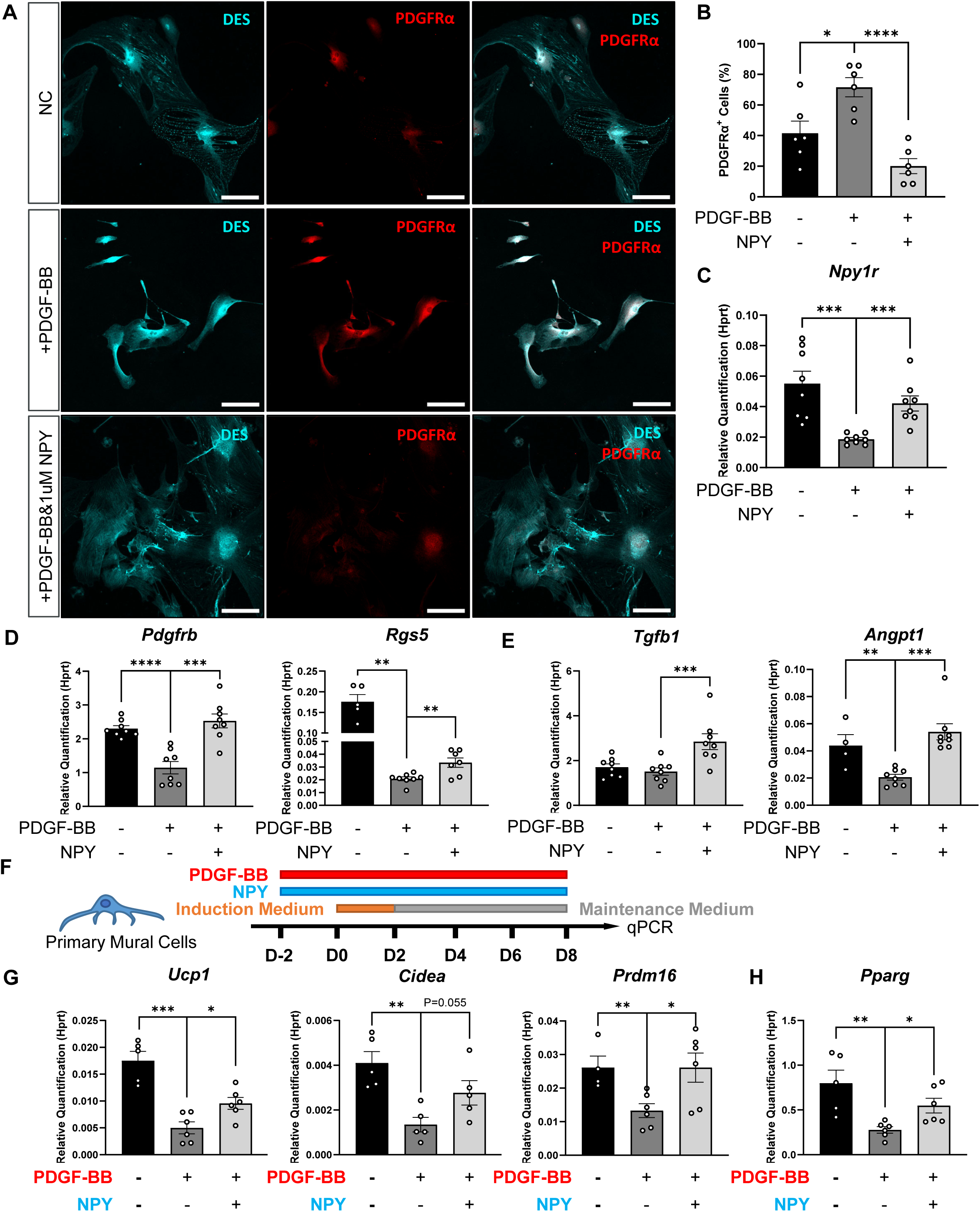
NPY sustains mural cells and their beiging ability against PDGF-BB. (A) Confocal images of primary mural cells stained with anti-DES (cyan) and anti-PDGFRα (red), cultured under different conditions. Scale bars=100μm. (B) The percentage of PDGFRα^+^ cells under different stimulating conditions (n=6). (C-E) The expression of (C) *Npy1r* (n=8), (D) mural cell markers *Pdgfrb* (n=8), and *Rgs5* (n=5, 8 & 8) and (E) secretory factors in maintaining vascular integrity including*Tgfb1* (n=8) and *Angpt1* (n=4&8&8) in primary mural cells cultured under different conditions. (F) Schematic of differentiating mural cells with PDGF-BB or PDGF-BB and NPY added (n=8). (G-H) The expression of (G) thermogenesis genes *Ucp1*, *Cidea*, *Prdm16*, and (H) beigeing gene *Pparg* in adipocytes differentiated from mural cells under different conditions indicated in the plots (n=5&6). The concentration of NPY is 1 μM and that of PDGF-BB is 250 ng/mL. All values are expressed as mean ± SEM, *p<0.05, **p<0.01, ***p<0.001, ****p<0.0001, Student T-tests.

**Supplementary Figure 9.**
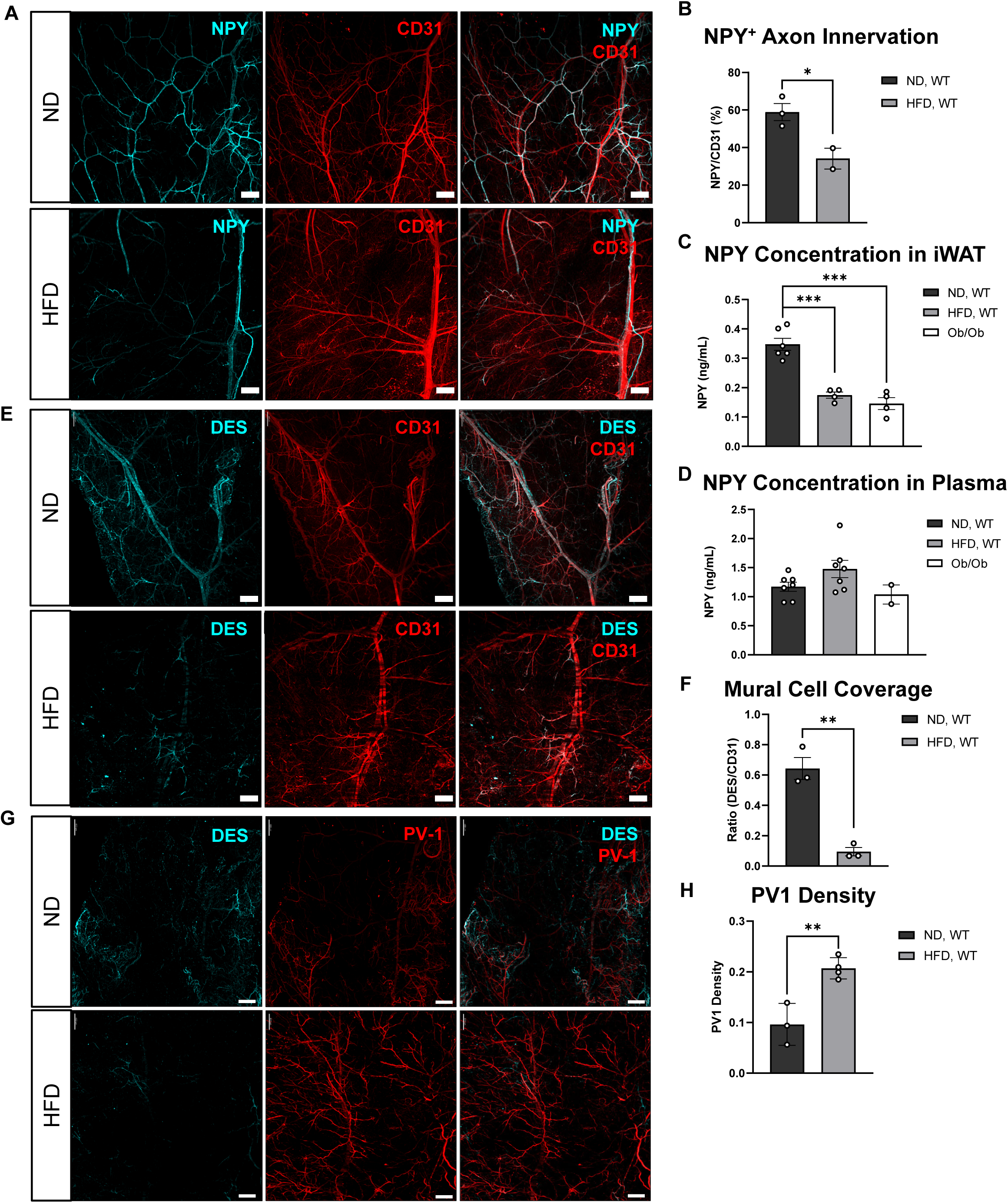
High-fat diet-induced obesity (DIO) depletes NPY^+^ innervation and mural cells and leads to vascular leakiness. (A) Light-sheet images of cleared iWAT of normal diet (ND)-treated (top) and high fat diet (HFD)-treated (bottom) 17-week-old male WT mice stained with anti-NPY (cyan) and anti-CD31 (red). Scale bars=500 μm. (B) NPY^+^ innervation calculated as the ratio of NPY to CD31 quantified based on images as in A (n=3&2). (C-D) The concentration of NPY in (C) iWAT (n=5, 4&4) and (D) blood plasma of ND and HFD-treated 17-week-old male WT mice, and ND-treated Ob/Ob 17-week-old male mice (n=6, 7&2). (E) Light-sheet images of cleared iWAT of ND– and HFD-treated 17-week-old male WT stained with anti-DES (cyan) and anti-CD31 (red). Scale bars=500 μm. (F) Mural cell coverage calculated as the ratio of DES^+^ cells to CD31^+^ cells quantified based on images as in E (n=3). (G) Light-sheet images of iWATs of ND– and HFD-treated 17-week-old male mice stained with anti-DES (cyan) and anti-PV1 (red). Scale bars=500 μm. (H) PV-1 density quantified based on images as in G (n=3&4). All values are expressed as mean ± SEM, *p<0.05, **p<0.01, ***p<0.001, ****p<0.0001, Student T-tests.

**Supplementary Figure 10.**
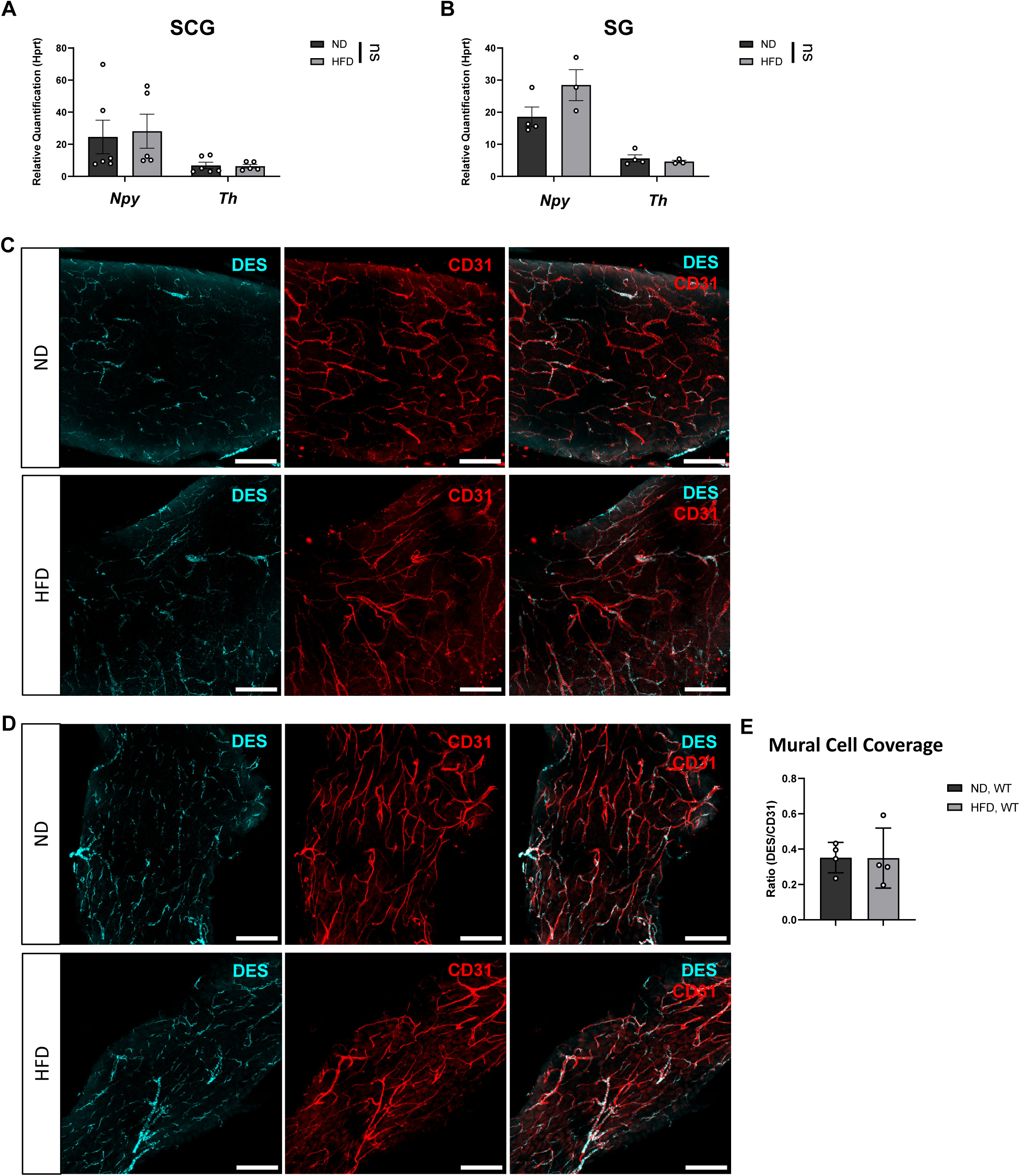
Obesity does not affect *Npy* expression in the ganglia. (A-B) The expression of *Npy* and *Th* in the (A) superior cervical ganglia (SCG, n=6&5) and (B) stellate ganglia (SG, n=4&3) of ND– and HFD-treated 17-week-old WT male mice. (C-D) The images of (C) cleared SCG and (D) cleared SG of ND– and HFD-treated 17-week-old WT male mice stained with anti-DES (cyan) and anti-CD31 (red) antibodies. Scale bars = 100 μm. (E) The quantification of mural cell coverage in SG (n=4). All values are expressed as mean ± SEM, *p<0.05, **p<0.01, ***p<0.001, ****p<0.0001, Student T-tests.

**Supplementary Figure 11.**
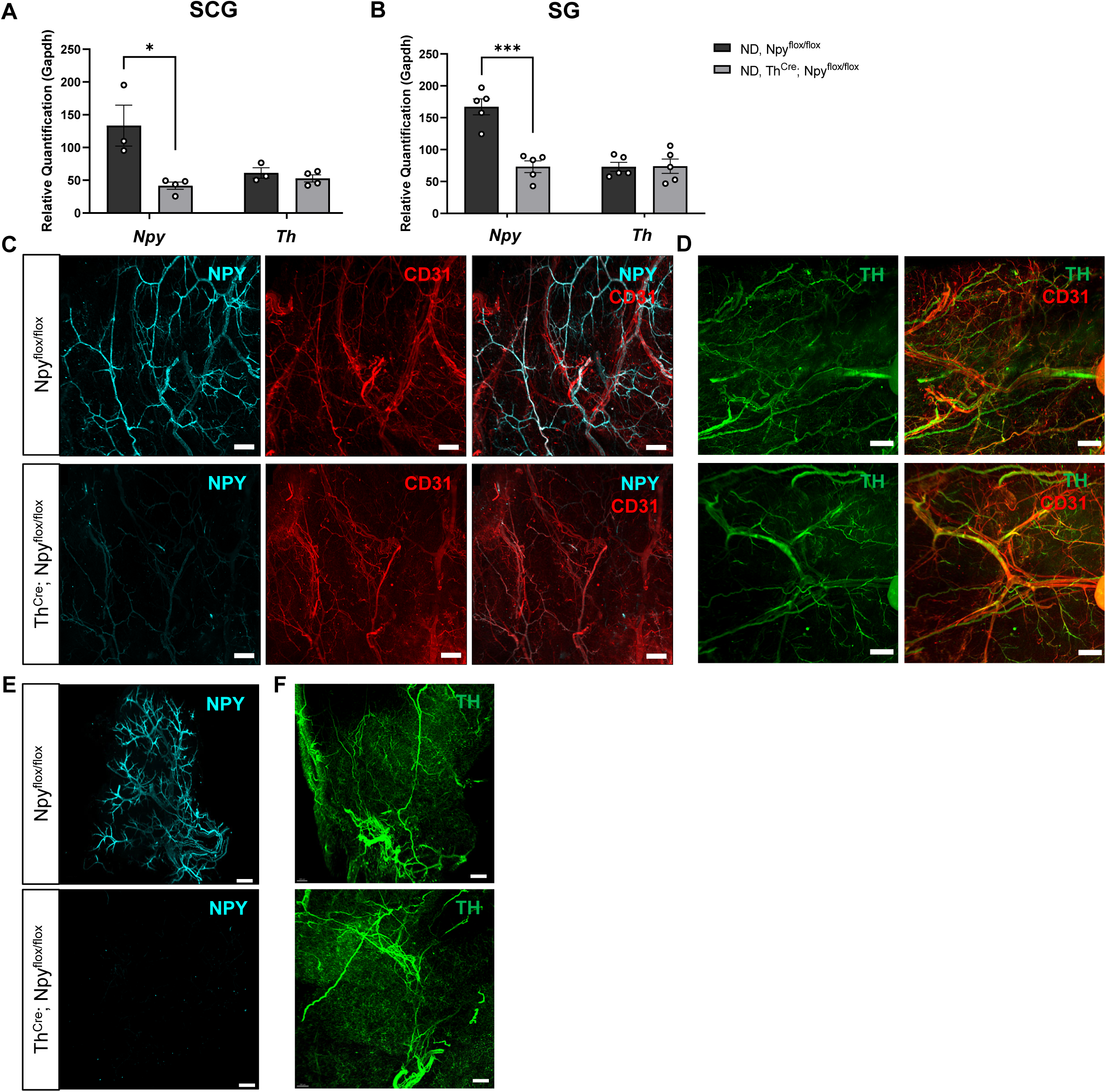
Th^Cre^; Npy^flox/flox^ mice deplete NPY from sympathetic nervous system without changing sympathetic innervation, central or plasma NPY level. (A-B) The expression of *Npy* and *Th* in (A) SCG (n=3&4) and (B) SG (n=5) of 12-week-old Th^Cre^; Npy^flox/flox^ mice and Npy^flox/flox^ male mice. (C-D) The image of cleared iWAT of 12-week-old Th^Cre^; Npy^flox/flox^ mice and Npy^flox/flox^ male mice stained with (C) anti-NPY (cyan) or (D) anti-TH (green) and anti-CD31 (red). Scale bar=500 μm. (E-F) Light-sheet image of cleared BAT of 12-week-old Th^Cre^; Npy^flox/flox^ mice and Npy^flox/flox^ male mice stained with (E) anti-NPY (cyan), scale bar= 500 μm, or (F) anti-TH (green), scale bar=200 μm. All values are expressed as mean ± SEM, *p<0.05, **p<0.01, ***p<0.001, ****p<0.0001, Student T-tests.

**Supplementary Figure 12.**
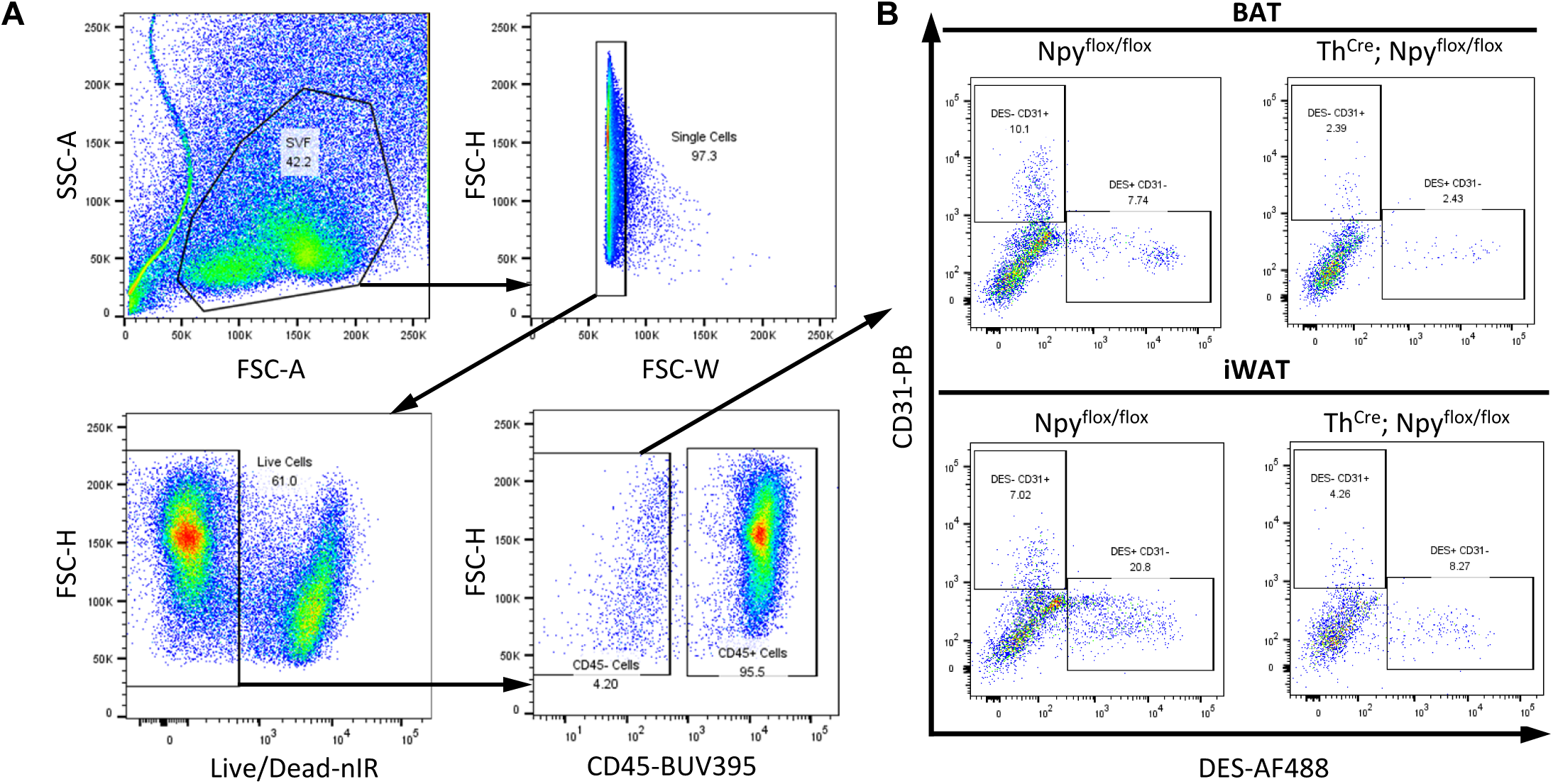
Loss of NPY from sympathetic neurons decreases the percentage of mural cells. (A) The gating strategy of flow cytometry to quantify the percentage of mural cells. (B) The representative flow cytometry analysis of the percentage of mural cells (DES^+^) in the BAT (top) and iWAT (bottom) of HFD-treated male Th^Cre^; Npy^flox/flox^ mice and Npy^flox/flox^ mice.

**Supplementary Figure 13.**
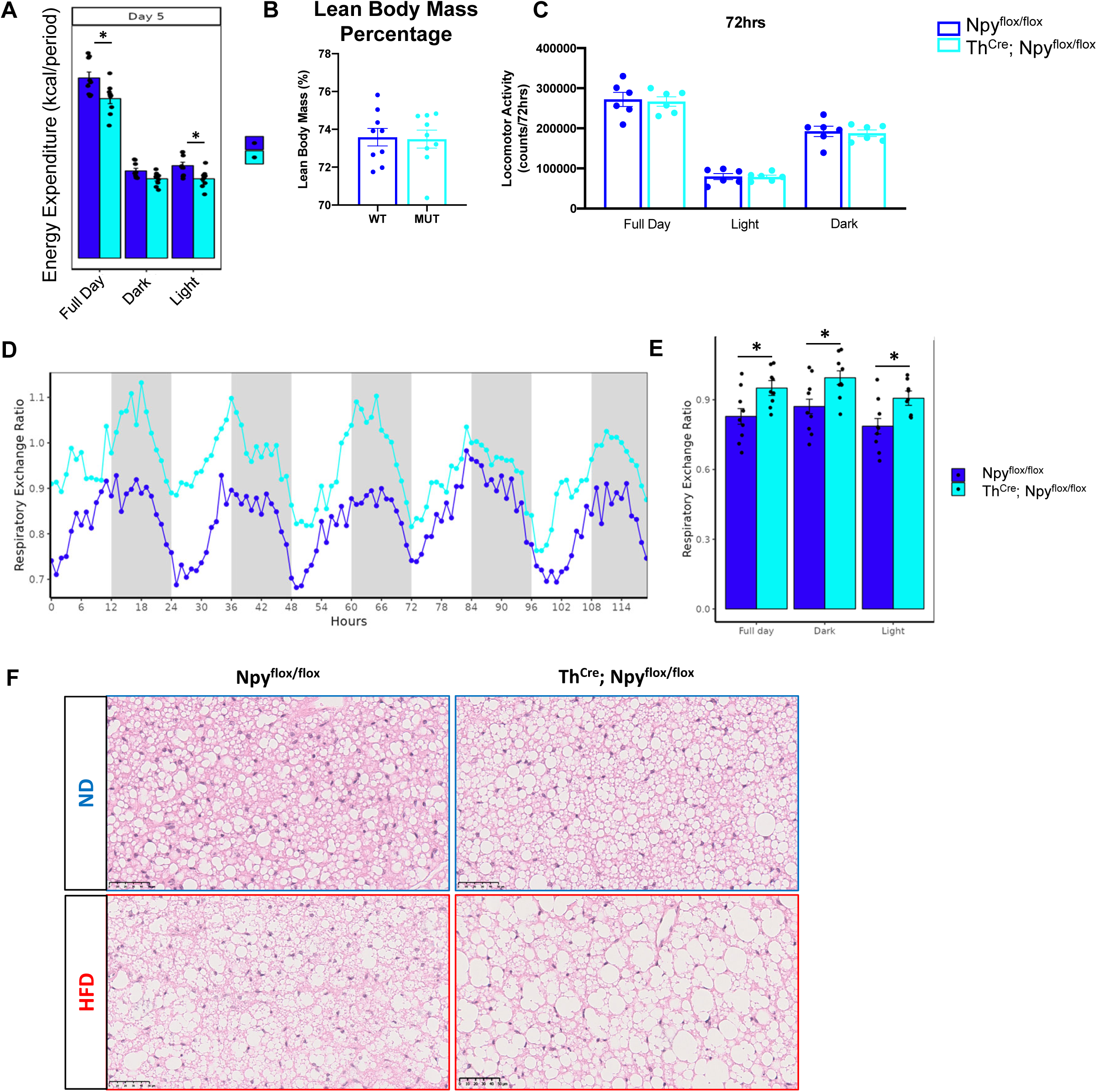
Loss of function of NPY from sympathetic neurons decreases energy expenditure, increases respiratory exchange ratio (RER) and lipid droplet size. (A) The quantification of daily energy expenditure recorded and quantified using an indirect calorimetry system (n=9). (B) The percentage of lean body weight (LBM) measured by MiniSpec MRI (n=9). (C) The locomotor activity measured using Multitake cage (n=6). (D) Respiratory exchange ratio (RER) recorded using a metabolic cage. (E) The quantification of B (n=9). (F) Histological staining of BATs of ND-(top) and HFD-(bottom) treated male Th^Cre^; Npy^flox/flox^ mice and Npy^flox/flox^ mice. Scale bars=50 μm.

**Supplementary Figure 14.**
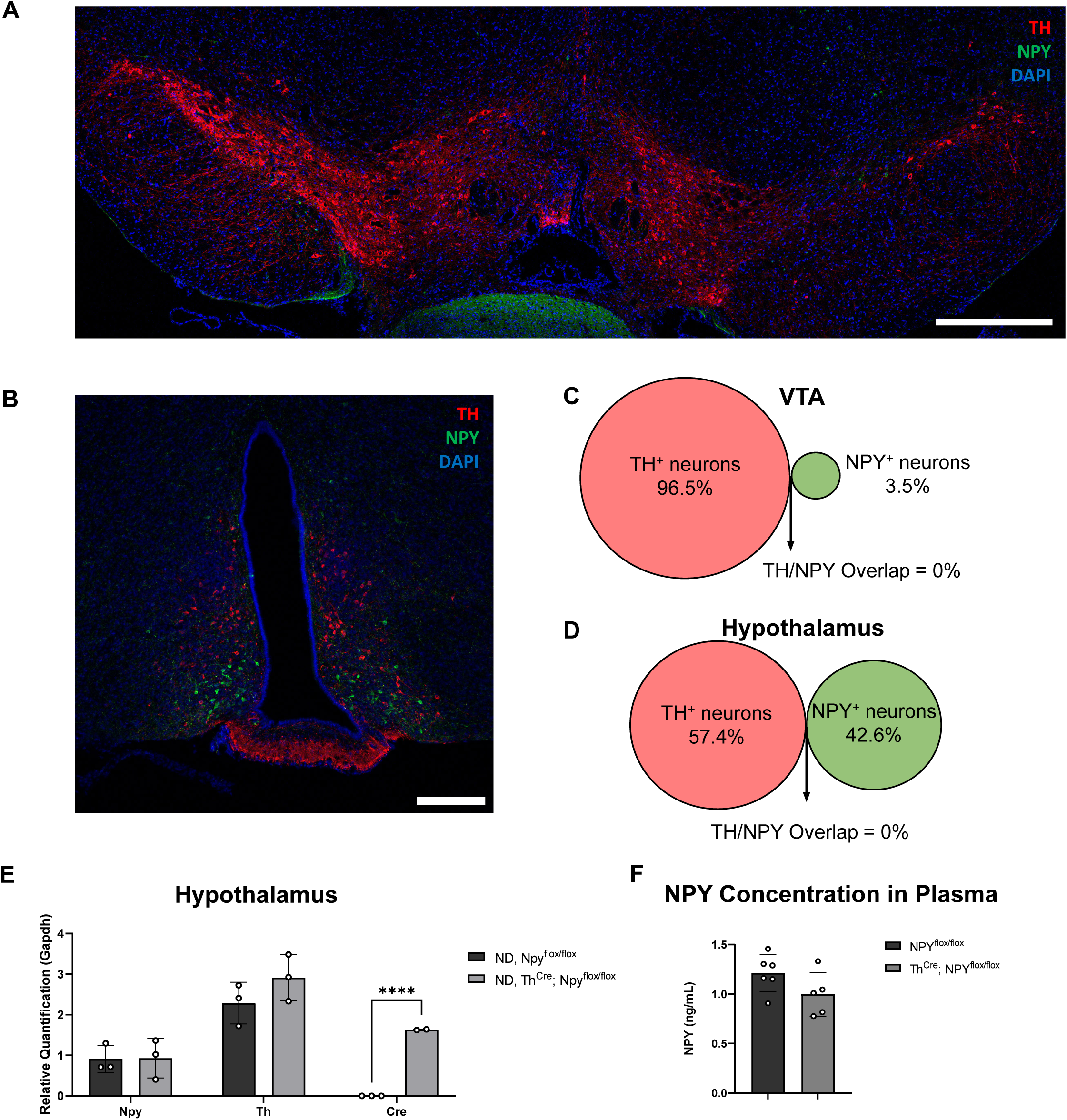
TH and NPY are not co-expressed in the VTA or hypothalamus, and Th^Cre^; Npy^flox/flox^ mice do not change Npy expression in hypothalamus or NPY concentration in plasma. (A) Confocal image showing a slice of the ventral tegmental area (VTA) of an 8-week-old NPY-GFP male mouse stained with anti-TH (red), anti-GFP (green), and DAPI (blue). Scale bar=500 μm (B) A confocal photomicrograph of a slice of arcuate nucleus of the hypothalamus of an 8-week-old NPY-GFP male mouse stained with anti-TH (red), anti-GFP (green), and DAPI (blue). Scale bar=200 μm (C-D) Quantification of TH/NPY-GFP overlap in (C) VTA (as in A, n=3) and (D) hypothalamus (as in B, n=3). (E) The expression of *Npy* (n=3)*, Th* (n=3), and *Cre* (n=2 & 3) in the hypothalamus of 12-week-old Th^Cre^; Npy^flox/flox^ mice and Npy^flox/flox^ male mice. (F) The concentration of NPY in the blood plasma of 12-week-old Th^Cre^; Npy^flox/flox^ mice and Npy^flox/flox^ male mice (n=6&5). All values are expressed as mean ± SEM, *p<0.05, **p<0.01, ***p<0.001, ****p<0.0001, Student T-tests.

**Supplementary Figure 15.**
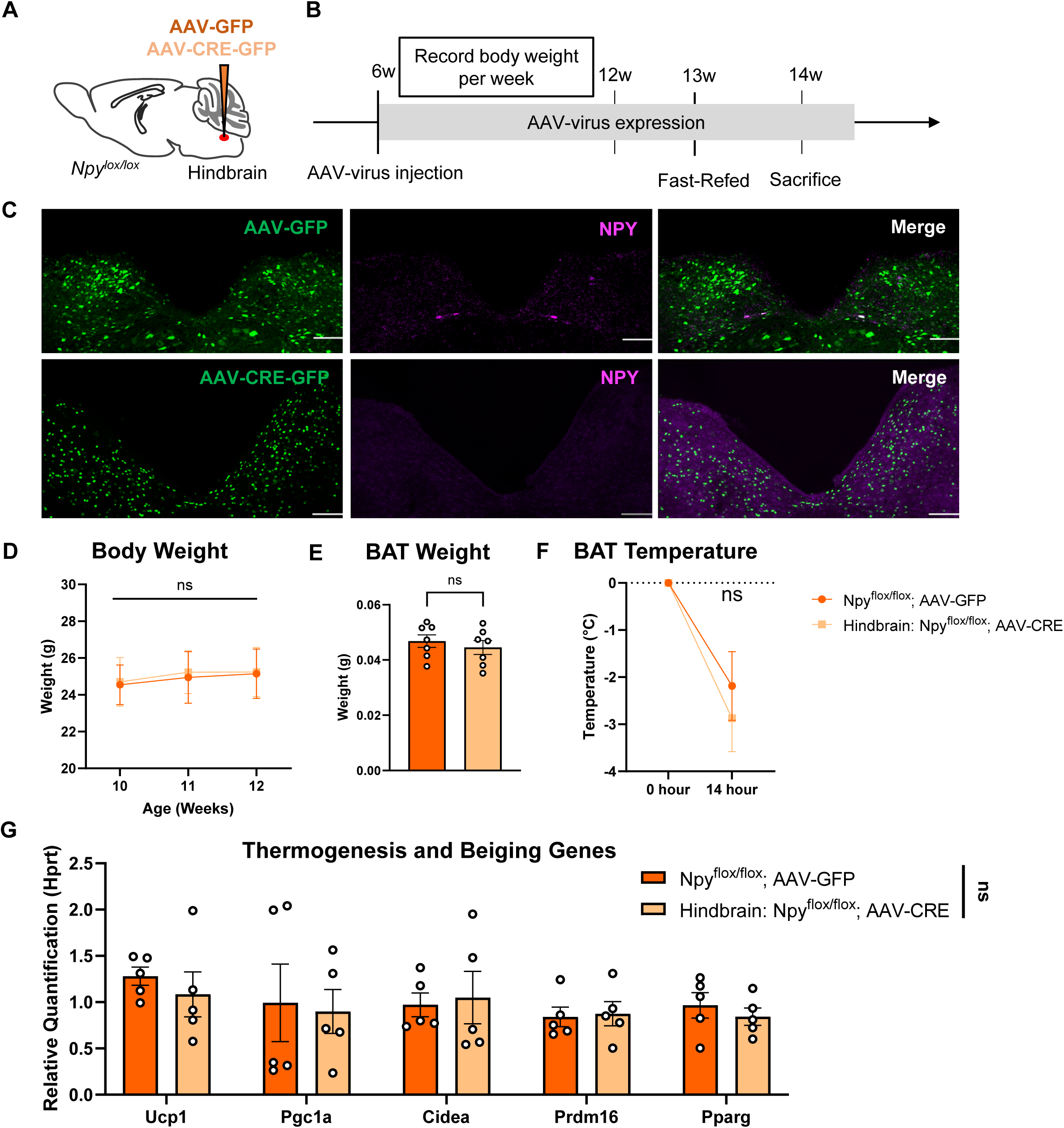
Ablating NPY from the hindbrain does not affect body weight, BAT weight or thermogenesis. (A) Schematic of knocking out NPY in the hindbrain by AAV injection. (B) The Schematic of metabolic phenotyping of mice with NPY knocked out in the hindbrain (Hindbrain: Npy^flox/flox^; AAV-CRE-GFP) and WT (Npy^flox/flox^; AAV-GFP) mice. (C) Confocal images of the sliced hindbrain of Hindbrain: Npy^flox/flox^; AAV-CRE-GFP and WT mice stained with anti-NPY (magenta). Scale bar=100 μm. (D) Weekly body weight of ND-treated male Hindbrain: Npy^flox/flox^; AAV-CRE-GFP and WT mice (n=7). (E) BAT weight of 14-week-old ND-treated male Hindbrain: Npy^flox/flox^; AAV-CRE-GFP and WT mice (n=7). (F) BAT temperature change after a 14-hour fast (n=7). (G) The expression levels of thermogenesis genes in the BAT of RT-housed ND-treated male Hindbrain: Npy^flox/flox^; AAV-CRE-GFP and WT mice (n=5).

**Supplementary Figure 16.**
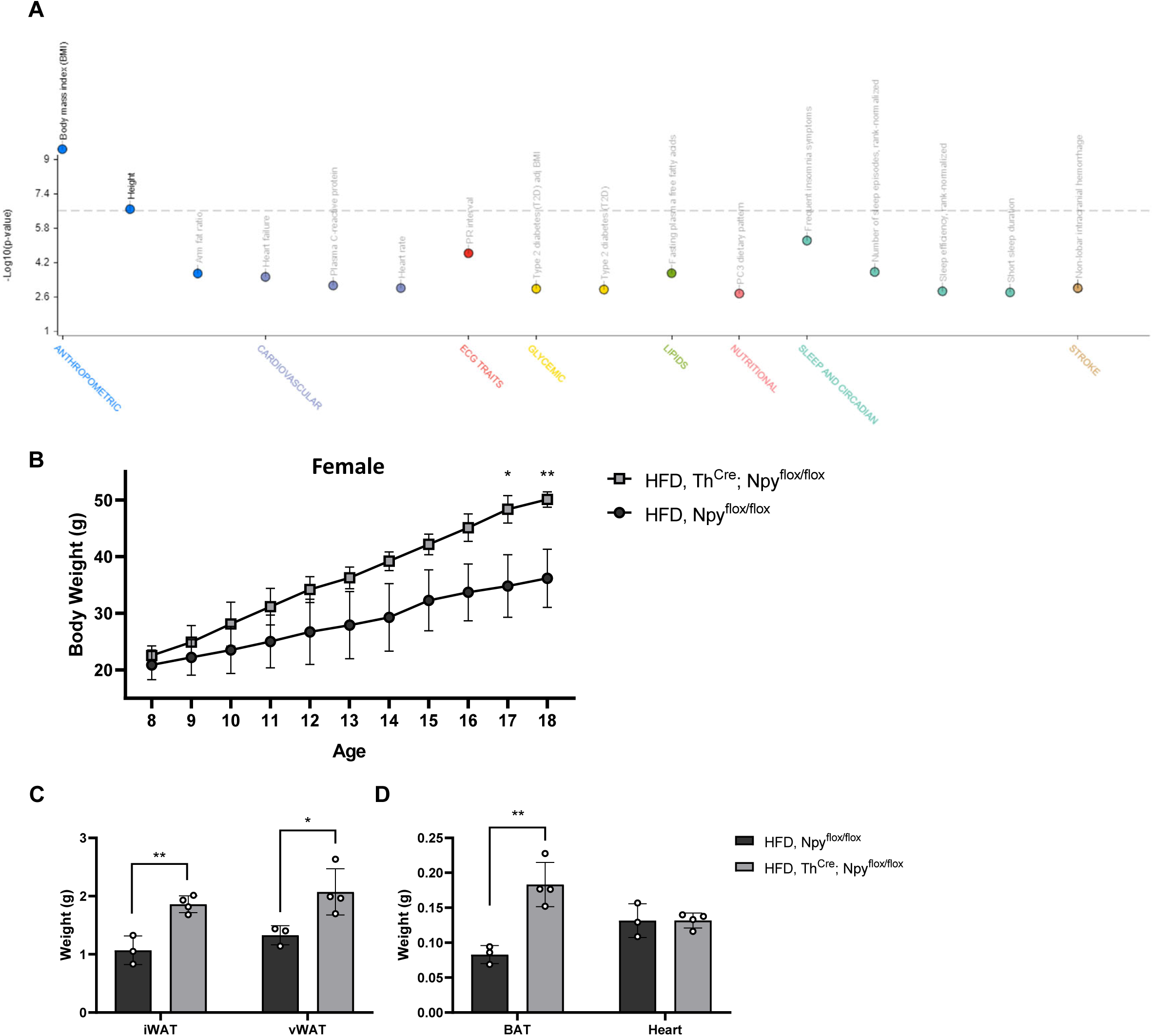
NPY variants significantly associate with human body mass index (BMI), and the metabolic phenotype of female Th^Cre^; Npy^flox/flox^ mice is similar to that of males. (A) Plot showing associations of human NPY gene with different traits^16^ (https://hugeamp.org/gene.html?gene=NPY). (B) The weekly body weights of female Th^Cre^; Npy^flox/flox^ mice and WT mice treated with high-fat diet (HFD) (n=4). (C-D) The weights of (C) iWAT and vWATs, and (D) BAT and hearts of HFD-treated 18-week-old female Th^Cre^; Npy^flox/flox^ mice and Npy^flox/flox^ mice (n=3&4). All values are expressed as mean ± SEM, *p<0.05, **p<0.01, ***p<0.001, ****p<0.0001, Student T-tests.

